# Stem cell models of TAFAZZIN deficiency reveal novel tissue-specific pathologies in Barth Syndrome

**DOI:** 10.1101/2024.04.28.591534

**Authors:** Olivia Sniezek Carney, Kodi William Harris, Yvonne Wohlfarter, Kyuna Lee, Grant Butschek, Arianna Anzmann, Steven M Claypool, Anne Hamacher-Brady, Markus Keller, Hilary J. Vernon

## Abstract

Barth syndrome (BTHS) is a rare mitochondrial disease caused by pathogenic variants in the gene TAFAZZIN, which leads to abnormal cardiolipin (CL) metabolism on the inner mitochondrial membrane. Although *TAFAZZIN* is ubiquitously expressed, BTHS involves a complex combination of tissue specific phenotypes including cardiomyopathy, neutropenia, skeletal myopathy, and growth delays, with a relatively minimal neurological burden. To understand both the developmental and functional effects of TAZ-deficiency in different tissues, we generated isogenic TAZ knockout (TAZ-KO) and WT cardiomyocytes (CMs) and neural progenitor cells (NPCs) from CRISPR-edited induced pluripotent stem cells (iPSCs). In TAZ-KO CMs we discovered evidence of dysregulated mitophagy including dysmorphic mitochondria and mitochondrial cristae, differential expression of key autophagy-associated genes, and an inability of TAZ-deficient CMs to properly initiate stress-induced mitophagy. In TAZ-deficient NPCs we identified novel phenotypes including a reduction in CIV abundance and CIV activity in the CIII2&CIV2 intermediate complex. Interestingly, while CL acyl chain manipulation was unable to alter mitophagy defects in TAZ-KO CMs, we found that linoleic acid or oleic acid supplementation was able to partially restore CIV abundance in TAZ-deficient NPCs. Taken together, our results have implications for understanding the tissue-specific pathology of BTHS and potential for tissue-specific therapeutic targeting. Moreover, our results highlight an emerging role for mitophagy in the cardiac pathophysiology of BTHS and reveal a potential neuron-specific bioenergetic phenotype.

## Introduction

Barth syndrome (BTHS, MIM#302060) is a rare, X-linked recessive genetic disorder that is clinically characterized by a cardiac phenotype of prenatal-onset of left ventricular noncompaction and infantile-onset cardiomyopathy, skeletal myopathy, growth delays, and intermittent neutropenia (1–4). There is a high risk of morbidity and mortality in BTHS, which is most often associated with cardiac disease (6–7). In contrast to many other mitochondrial disorders, BTHS patients have a minimal neurological burden, with a mild neurodevelopmental profile reported in some patients (1,5).

BTHS is caused by pathogenic variants in the gene *TAFAZZIN (TAZ,* MIM#300394), located on chromosome Xq28 (8–9). The encoded protein, TAFAZZIN (TAZ), is a ubiquitously expressed mitochondrial transacylase that is critical for the remodeling of the phospholipid cardiolipin (CL), which is a unique component of inner mitochondrial membranes (IMM) (8, 10–11). CL remodeling by TAZ yields a tissue-specific acyl chain pattern, with heart and muscle CL acyl chains predominantly composed of linoleic acid (LA) acyl moieties (>80%) and brain tissue acyl chains composed of longer chain fatty acids and a higher oleic acid (OA) content (12–15). Deficiency in TAZ-based CL remodeling results in a reduction in remodeled (mature) CL, an increase in the unmodeled intermediate monolysoCL (MLCL), and an abnormal acyl chain composition on the remaining CL species, shifted towards increased saturation (1,16). The primary diagnostic metabolic defect in BTHS is an elevated MLCL:CL ratio, which is detected in 100% of affected individuals and every TAZ-deficient model described to date (17–22). A recent study established that the major determinants of CL acyl chain content are fatty acid pools of individual cell and tissue types (13). Importantly, the cellular CL content and composition can be manipulated *in vitro* by nutritional supplementation of specific fatty acids (13).

Despite knowledge of the genetic basis and primary biochemical abnormalities in BTHS, it remains unclear how abnormal CL content contributes to tissue specific cellular pathologies (23–27). Prior studies in cell and animal models of BTHS have identified mitochondrial respiratory chain dysfunction, including oxidative phosphorylation (OXPHOS) and bioenergetic defects, supercomplex destabilization, and reduced complex I (CI) expression and enzymatic function (17, 28–30).

Emerging evidence suggests a novel role for dysfunctional mitochondrial quality control mechanisms, specifically mitophagy, in the pathogenesis of BTHS (17, 31–35). Mitophagy, the targeted, sequestration and degradation of damaged mitochondria in autophagosomes, is an essential cellular process that, when defective, leads to the accumulation of dysfunctional mitochondria and increased cellular oxidative stress (36–38). Mitophagy can be initiated by several cellular cascades and CL has known roles in receptor-mediated pathways where it facilitates binding of LC3, a key autophagosomal membrane component that recognizes mitochondrial outer membranes marked for degradation (39–41). Conversely, MLCL, which is accumulated in BTHS, is unable to properly bind and recognize LC3, likely contributing to failure of autophagosomes to properly localize (41–42). Mitophagy represents an enticing candidate for study in BTHS, as impaired mitophagy has been implicated in the pathogenesis of related cardiac complications including dilated and hypertrophic cardiomyopathy, ischemia-reperfusion injury, and diabetic cardiomyopathy (37). In addition, mitophagy is an essential component of cardiac differentiation, maturation, and survival, and, when defective, contributes to abnormal cardiac morphogenesis and chamber formation (36, 43–44).

We generated and characterized a novel, TAFAZZIN-knock out (TAZ-KO) induced pluripotent stem cell (iPSC) model and differentiated cells into an affected cell type in BTHS (cardiomyocytes, CMs) and a relatively spared cell type (neural progenitor cells, NPCs) in order to understand and manipulate tissue-specific mechanisms of mitochondrial dysfunction. We identified important mechanisms of cell type-specific mitochondrial dysfunction including dysregulation of mitophagy, which was more apparent in CMs compared to NPCs. We also identified cell type-specific respiratory chain assembly and function abnormalities, with defective intermediate complex assembly of CIII/CIV species apparent in TAZ-deficient NPCs but not CMs. To test whether manipulation of CL acyl content could modify these phenotypes, we employed a nutritional manipulation strategy to alter CL acyl content and saturation. Interestingly, while we were able to predictably alter CL acyl chain content, there were few observable effects on mitochondrial dysfunction. Altogether, our findings suggest that TAFAZZIN- deficiency and abnormal CL content havs tissue-specific effects on mitochondrial dysfunction, with a mitophagy defect contributing to CM pathology and Complex III/IV assembly contributing to possible neuronal pathology. Our results also suggest that tissue-specific therapeutic approaches, beyond CL side chain modification, may be necessary to treat the pleiotropic effects of BTHS.

## Results

### Generation and phenotyping of a novel TAFAZZIN-deficient iPSC model

We generated a novel *TAFAZZIN* knock-out (TAZ-KO) induced pluripotent stem cell (iPSC) model using CRISPR/Cas9 genome editing in a wildtype (WT) male iPSC line using the Moyer and Holland Protocol (45). Using two single guide RNAs (sgRNAs) targeting the 3’-end of exon 2 of the *TAFAZZIN* gene, we generated and isolated four clones with a 50 bp deletion, c.456_505del, encompassing a predicted acyltransferase domain (Fig. S1 A). This region also coincides with a region of *TAFAZZIN* where multiple pathological variants have been identified in affected individuals (Fig. S1 B) (10). The 50 bp deletion was confirmed by qRT-PCR, and mRNA expression was detected at a similar abundance to WT, but at a smaller size, reflective of the deletion (Fig. 1A). Immunoblotting of four individual edited clones demonstrated undetectable levels of TAZ protein (Fig 1B).

**Fig. 1.**
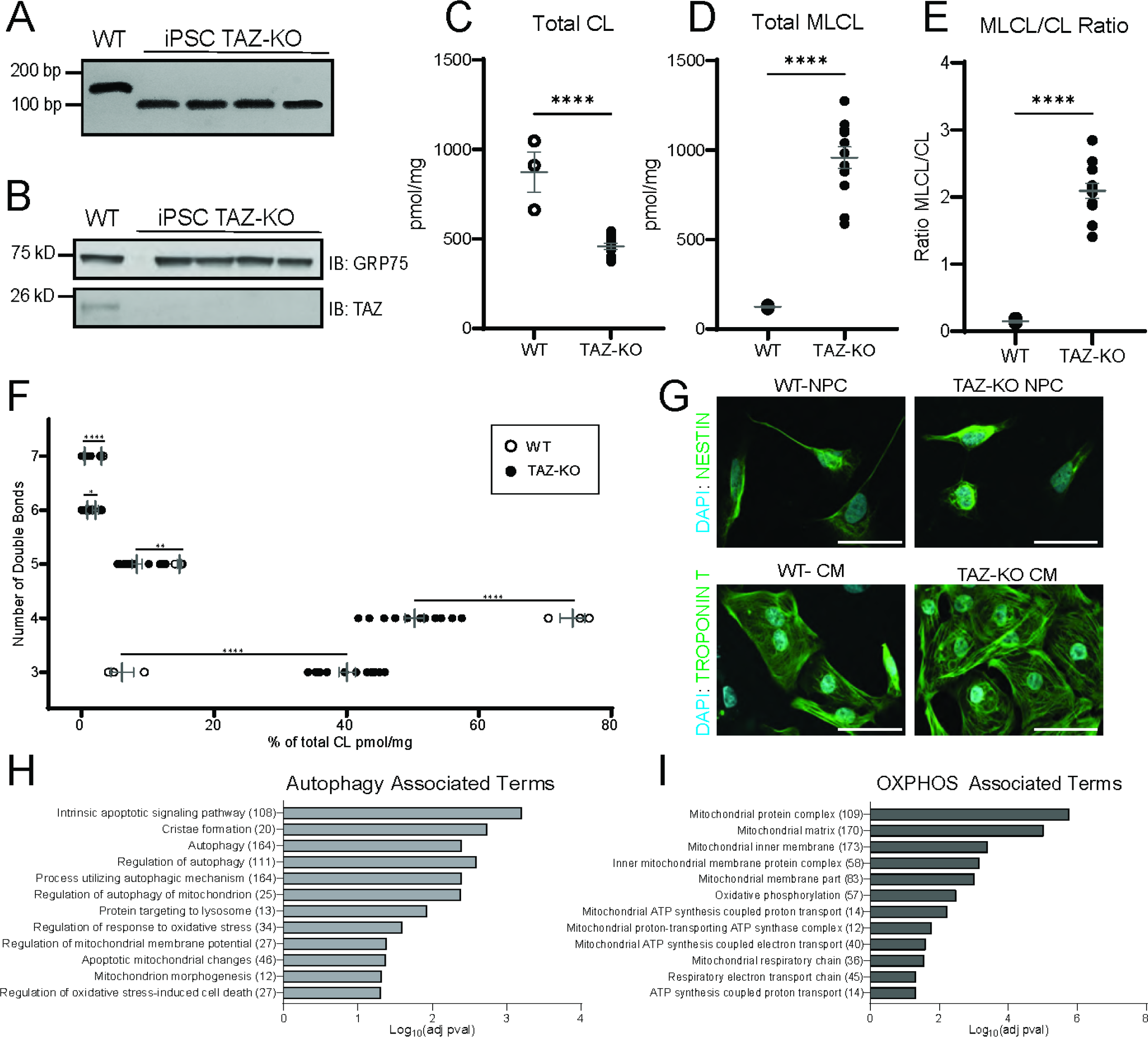
***TAZ-deficient iPSC phenotyping, differentiation, and RNAseq analysis.*** (A) RT-PCR of mRNA extracted from undifferentiated (UD) WT and TAZ-KO iPSCs using primers to capture the edited region. (B) Whole cell lysate (60 µg) of indicated cell lines were immunoblotted for TAZ, and GRP75 as loading control. (C-E)The abundance of CL and MLCL were determined via shotgun lipidomics mass spectrometry and the MLCL/CL ratio is reported for each line with values normalized to protein content. (F) The distribution of CL species containing specified numbers of double bonds is reported as percentage of total CL. (G) Differentiation of iPSCs into neural progenitor cells (NPCs) and cardiomyocytes (CMs) was verified via imunofluorescence for specified lineage markers. Shown are representative images for each cell type and genotype. Scale bar, 100 µm. (H and I) RNAseq was performed in triplicate on WT an TAZ-KO UD, CM, and NPCs. Differentially expressed genes were analyzed for gene ontology (GO) Terms. Terms statistically significantly lower in TAZ- KO UD/WT are reported for authophagy associated terms, as well as oxidative phosphorylation associated terms. Adjusted p-values <0.05. Significant differences are indicated; *≤0.05, **≤0.005, ***≤0.0005, and ****≤0.00005.

Shotgun lipidomics mass spectrometry analysis of CL and MLCL species revealed significantly decreased CL in TAZ-KO iPSCs (p<1x10^-4^), significantly increased MLCL (p<1x10^-4^), and a significant increase in the MLCL/CL ratio (p<1x10^-4^) (Fig. 1 C-E). Characteristic of lack of TAFAZZIN-based remodeling, CLs in TAZ-KO iPSCs were more saturated, with significantly increased 3 double bond containing species (p<1x10^-6^) and significantly decreased 4-7 double bond containing species (p= 3x10^-4^, p= 1.4x10^-3^, p= 0.0293, p< 1x10^-6^, respectively) (Fig. 1 F). Taken together, our TAZ-KO iPSC model recapitulates the pathognomonic biochemical defect in BTHS patients and other published *TAFAZZIN-*deficient cell models (17–22), thus validating its use as a BTHS model.

### Differentiation of TAZ-KO iPSCs into clinically relevant cell types and identification of dysregulated pathways

We performed RNAseq (n=3 biological replicates) of undifferentiated (UD) WT and TAZ-KO iPSCs. Analysis of differentially expressed genes for Gene Ontology (GO) terms demonstrated significant enrichment for pathways involved in mitochondrial quality control (MQC) including, regulation of autophagy (P=0.000000989), intrinsic apoptotic signaling pathway (P=0.00000255), intrinsic apoptotic signaling pathway in response to endoplasmic reticulum stress (P=0.0000026) and macroautophagy (P=0.00000385) (Fig. 1H). In addition to MQC pathways, TAZ-KO UD iPSCs demonstrated an enrichment for genes associated with OXPHOS, including Oxidative phosphorylation, inner mitochondrial membrane protein complexes, and ATP synthesis coupled electron transport (Fig. 1I).

To study tissue-specific phenotypes in BTHS, we differentiated WT and TAZ-KO iPSCs into CMs and NPCs, representing cell types that are affected (CMs) vs relatively spared (NPCs) in BTHS. qRT-PCR of cell type-specific markers confirmed presence of pluripotency markers (NANOG and OCT4) in undifferentiated iPSCs and absence of these markers in differentiated cell types, as well the gain of neural-specific marker (PAX6) and CM-specific marker (MHC6), respectively (Fig. S2C). We further investigated cell type-specific markers via immunofluorescence staining of nestin in NPCs and cardiac troponin T (CTNN1) in CMs (Fig. 1G).

### TAZ-KO iPSC CMs have abnormal mitochondrial morphology and ultrastructure

We followed up potential consequences of the disturbed MQC-related pathways identified in UD iPSCs by examining mitochondrial morphology in differentiated cell types, WT and TAZ-KO iPSC CMs and NPCs, via immunofluorescence staining of the outer mitochondrial membrane marker TOM20 (Fig. 2 A&C). We analyzed mitochondrial networks using the ImageJ/Fiji plugin Mitochondrial Analyzer (48). Compared to WT CMs, mitochondrial networks in TAZ-KO CMs displayed a more fragmented morphology, with significantly fewer, smaller, and less branched mitochondria (Fig. 2B). Interestingly, this phenotype was specific to TAZ-deficient CMs, as mitochondrial morphological parameters were less disturbed in TAZ-KO NPCs when compared to WT-NPCs (Fig. 2C-D).

**Fig. 2.**
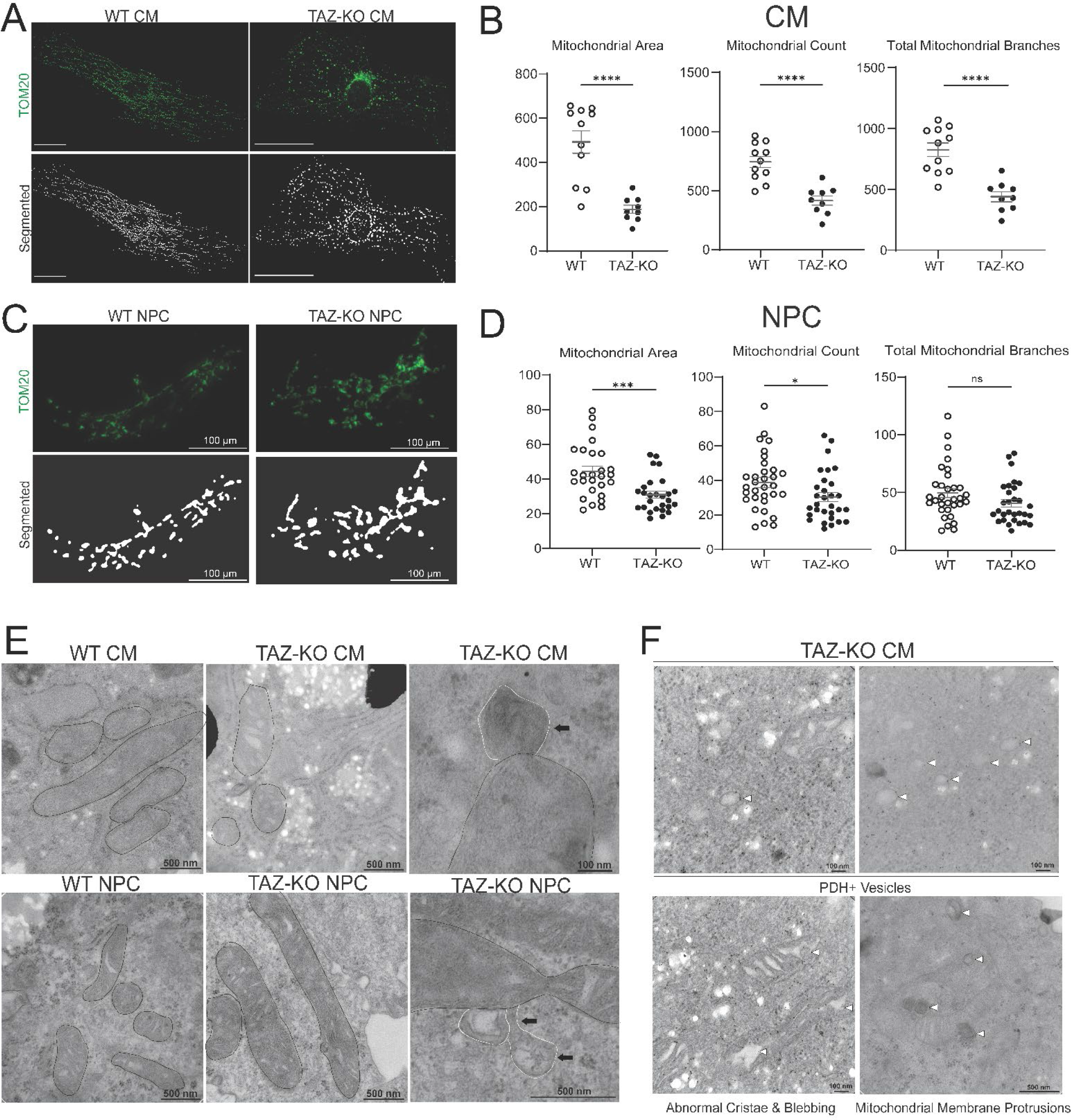
***TAZ-deficient CMs demonstrate aberrant mitochondrial morphology and ultrastructure.*** (A) Representative immunofluorescence images of WT and TAZ-KO CM TOM20 stained mitochondria (upper panel) and image segmentation (lower panel). Scale bars, 100 µm. (B) Mitochondrial networks from individual cells were quantified using Mitochondrial Analyzer R package and metrics from individual cells are reported as well as the mean, and standard error of the mean. (C) Representative immunofluorescence images of WT and TAZ-KO NPC TOM20 stained mitochondria (upper panel) and image segmentation (lower panel). (D) Mitochondrial networks from individual cells are reported as well as the mean, and standard error of the mean. (E) Transmission electron microscopy was performed on CMs and NPCs of both genotypes. Individual mitochondria are outlined in black, mitochondrial membrane protrusions are outlined in white. (F) Immunoelectron microscopy of immunogold-stained PDH in TAZ-KO CMs is shown with white arrows indicating PDH+ vesicular structures. Significant differences are indicated; *≤0.05, **≤0.005, ***≤0.0005, and ****≤0.00005.

We then characterized mitochondrial ultrastructure with transmission electron microscopy (TEM) of WT and TAZ-KO CMs and NPCs. Compared to WT, TAZ-KO CMs contained fewer mitochondria and less organized cristae, whereas TAZ-KO NPC mitochondrial ultrastructure was similar to WT NPCs (Fig. 2E). Interestingly, both TAZ-KO CMs and NPCs showed an increase in mitochondrial membrane protrusions, suggestive of mitochondrial derived vesicles (MDVs) (62, 64) (Fig. 2E). Immunogold staining revealed the presence of PDH-positive vesicles in the cytosol of TAZ-KO, but not WT CMs (Fig. 2F), in further support of the activation of MDV pathways in TAZ-KO CMs.

Interestingly, MDVs can function in MQC and can compensate for deficiencies in canonical, autophagosome-mediated mitophagy (65). To characterize levels of canonical mitophagy in TAZ-KO CMs and NPCs, at baseline and after mitophagy induction with the depolarizing agent carbonyl cyanide m-chlorophenylhydrazine (CCCP), we utilized a fluorescence Dojindo Mtphagy Dye-based assay that allows for the detection of mitochondrial delivery to lysosomes in *live* cells (63). As expected, WT CMs and NPCs showed increased levels of mitophagy induction upon CCCP treatment, reflected by the increased colocalization of mitochondrial-localized Mtphagy Dye with lysosome-localized Lyso Dye (Fig. 3 A-D). Notably, such CCCP-induced mitophagy response was absent in TAZ-KO CMs. Mitophagy levels in TAZ-KO NPCs were responsive to CCCP treatment, albeit overall reduced compared to WT NPCs.

**Fig. 3.**
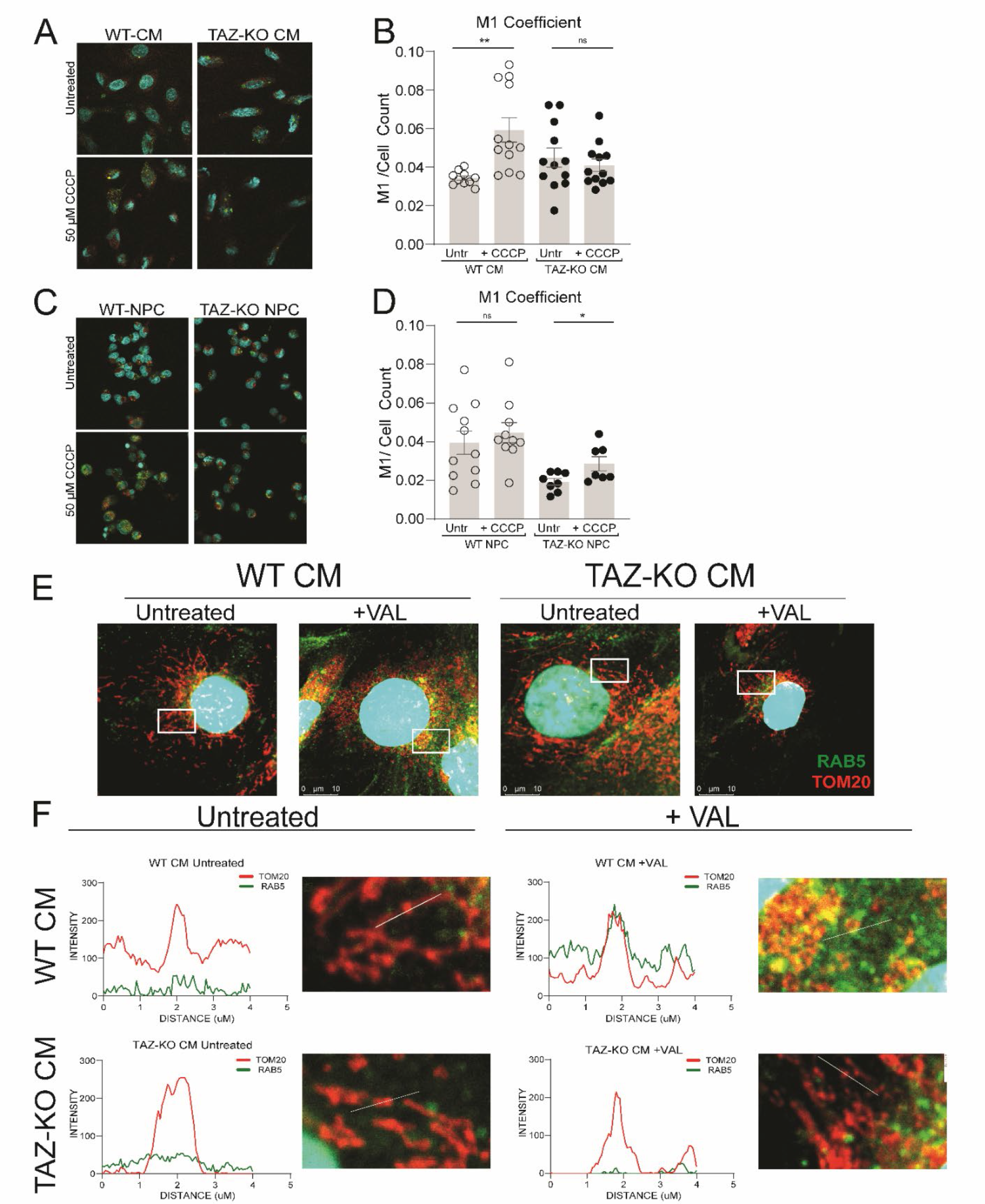
***TAZ-KO iPSC-CMs are defective mitophagy initiation and display inability to recruit mitophagy proteins to the mitochondria.*** (A) WT and TAZ-KO CMs, either untreated, or treated with 50 µM CCCP for 30 minutes, were co- stained with DAPI, Dojindo Mtphagy Dye and Lyso Dye. Scale bar, 50 µm. (B) M1 Coefficient of correlation between Dojindo Mtphagy Dye and Lyso Dye was measured using the Fiji plugin ‘Coloc 2’ and normalized to cell count. (C) Representative immunofluorescence images of WT and TAZ-KO NPCs, either untreated or treated with 50 µM CCCP for 30 minutes, stained with Dojindo Mtphagy dye. (D) M1 coefficient of correlation was measured using Fiji and reported normalized to cell count. (E) Representative images of WT and TAZ-KO CMs untreated or treated with VAL for 6 hours were stained with RAB5A (green) and TOM20 (red). (F) Intensity plots of RAB5A (green) and TOM20 (red) were used to measure colocalization of the two signal intensities from regions defined by the white boxes in panel E. Significant differences are indicated: *≤0.05, **≤0.005, ***≤0.0005, and ****≤0.00005

We thus next tested whether non-canonical MQC pathways might be enhanced in TAZ-KO CMs that could support compensation for impaired canonical mitophagy in these cells. To that end, we evaluated mitochondrial recruitment of RAB5, an early endosomal small GTPase that can mediate alternative mitochondrial degradative processing via endosomal uptake of mitochondria (66), and via entrance of endosomes into intra-mitochondrial compartments (67). In response to mitochondrial depolarization with the potassium ionophore valinomycin, WT-CMs demonstrated robust recruitment of RAB5 to the mitochondria, while activation of mitochondrial targeting by RAB5 was found to be less pronounced TAZ-KO CMs (Fig. 3 E-F). Together, these findings support specific deficiencies in canonical and non-canonical MQC pathways in TAZ-KO cells that appear to manifest more severely in TAZ-KO CMs vs. NPCs and are associated with potentially compensatory upregulation of MDVs.

### Cell type specific characterization of oxidative phosphorylation (OXPHOS) complex assembly and activity in iPSC differentiated cell types

Various respiratory chain complex assembly defects have been reported in TAZ-deficient models, including supercomplex destabilization (17, 28–30). Thus, we characterized OXPHOS complex assembly and activity via blue native polyacrylamide gel electrophoresis (Blue Native PAGE) and immunoblotted for complexes I, II, III, and IV alone or in combination (Total OXPHOS). In both WT and TAZ-KO genotypes, we identified cell type-specific patterning of OXPHOS supercomplex and intermediate complex assembly and activity (Fig. 4 A-B, C-D), with iPSC-derived cardiomyocytes having greater expression and abundance of all OXPHOS intermediate and supercomplexes compared to NPCs and UDs (Fig. 4 A-B, D-E). Given substantial differences in respiratory capacity and substrate preferences across tissues, these changes are consistent with the observations that all respiratory complexes, except CII, are higher in cardiac cells, which are more energy dependent (49–50). In addition, we evaluated CI and CIV in gel activity and observed higher CI and CIV activity, especially CI and CIV containing supercomplexes, in CMs compared to other cell types (Fig. 4 C & F). This was agnostic to genotype, with both WT and TAZ-KO cells showing the same cell type-specific assembly patterns.

**Fig. 4.**
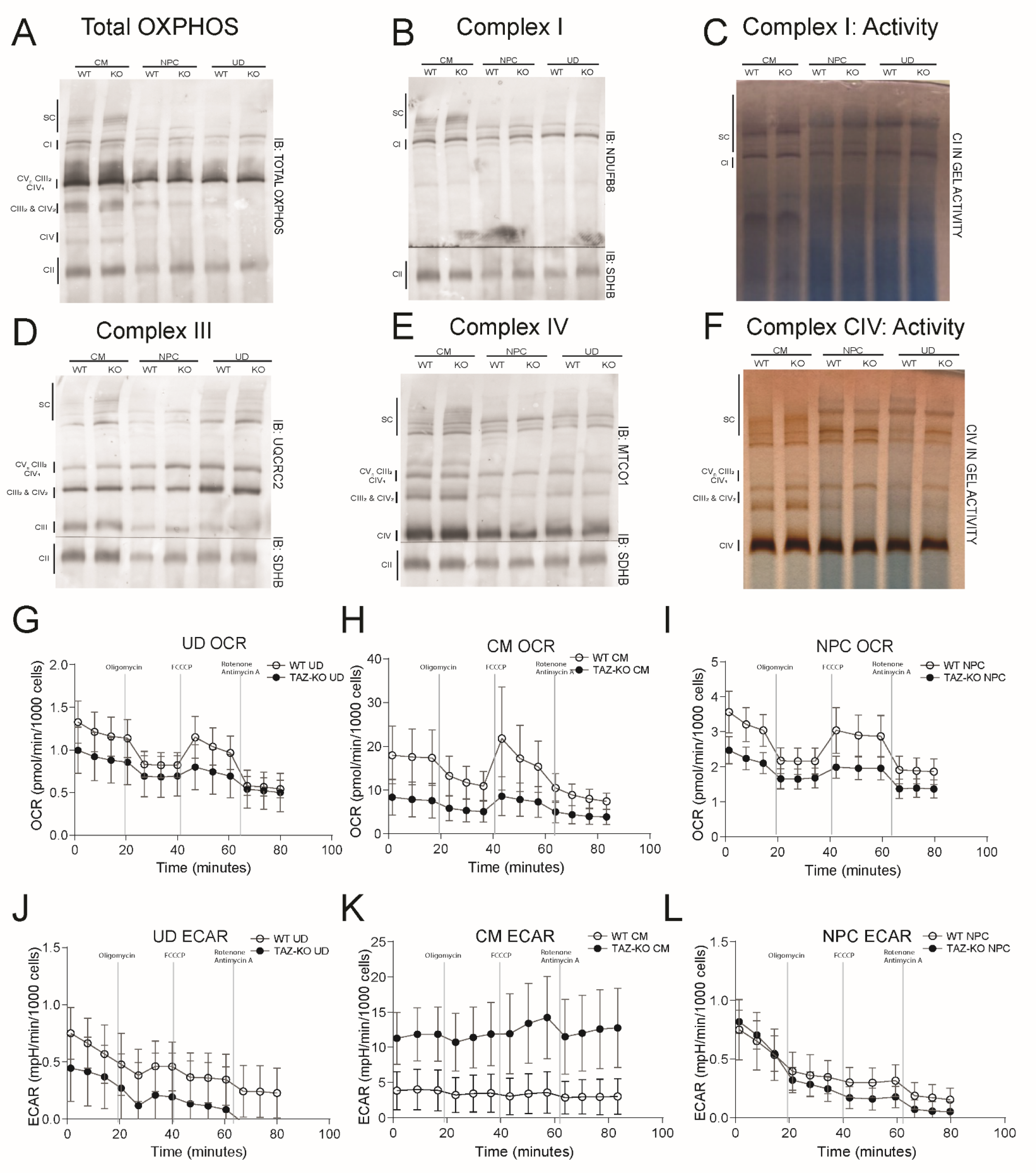
***TAZ-KO CMs and NPCs show disparate defects in OXPHOS assembly and function.*** (A) Cell pellets of 1x10^6^ cells were resolved using blue native poly acrylamide gel electrophoresis (Blue Native- PAGE) and immunoblotting was performed to assess (A) total OXPHOS assembly (B) Complex I and II, (D) Complex III and II, and (E) Complex IV and II. In gel activity was performed using native-page electrophoresis on 1x10^6^ cells for (C) Complex I and (F) Complex IV. (G-I) Oxygen consumption was measured using a seahorse extracellular flux analyzer (XFe96) for indicated cell types and genotypes and is normalized per 1000 cells. (J-L) Extracellular acidification rate is reported per 1000 cells.

Interestingly, we observed a reduction in CIV abundance and CIV activity in the CIII2&CIV2 intermediate complex in TAZ-deficient NPCs (Fig. 4 E-F). CIII2&CIV2 abundance was measured as a ratio to CII revealing a ratio of 0.55 in WT NPCs and 0.34 in TAZ-KO NPCs reflecting a nearly 40% reduction. This reduction was specific to CIV, as we observed no reduction in the CIII portion of the complex (Fig. 4D).

To characterize overall changes in mitochondrial respiratory capacity, we evaluated mitochondrial oxygen consumption (OCR) in WT and TAZ-KO iPSC CMs, NPCs, and UDs, and revealed a reduction in OCR, and maximal respiratory capacity, in all TAZ-KO cell types compared to WT counterparts, with the most significant changes in TAZ-KO CMs (Fig. 4 G-I). In addition, TAZ-KO CMs showed significantly reduced basal respiration, ATP production, and spare respiratory capacity (Fig. S3). The extracellular acidification rate (ECAR) was notably higher in TAZ-KO CMs indicating elevated glycolytic activity compared to WT counterparts (Fig. 4K). ECAR levels in TAZ-KO UDs and NPCs were less significantly different compared to WT (Fig. 4 J and K).

### Cell type specific analysis of CL content and alteration of CL acyl composition and saturation

To elucidate the role of the CL acyl complement on downstream cellular dysfunction in TAZ-deficiency, we subjected cells to nutritional supplementation with either 100 µM LA or OA for 8 days and assessed CL species and saturation. As expected, at baseline, TAZ-KO UDs, CMs, and NPCs had increased CL saturation (Fig. 5 A) and increased species containing fewer double bonds (Fig. 5 D-F) compared to WT counterparts. Characteristic of lack of TAZ-based remodeling, TAZ-KO cells showed lower CL abundance (Fig. 5 G) and an elevated MLCL/CL ratios both pre-supplementation and post-supplementation (Fig. 5 H).

**Fig. 5.**
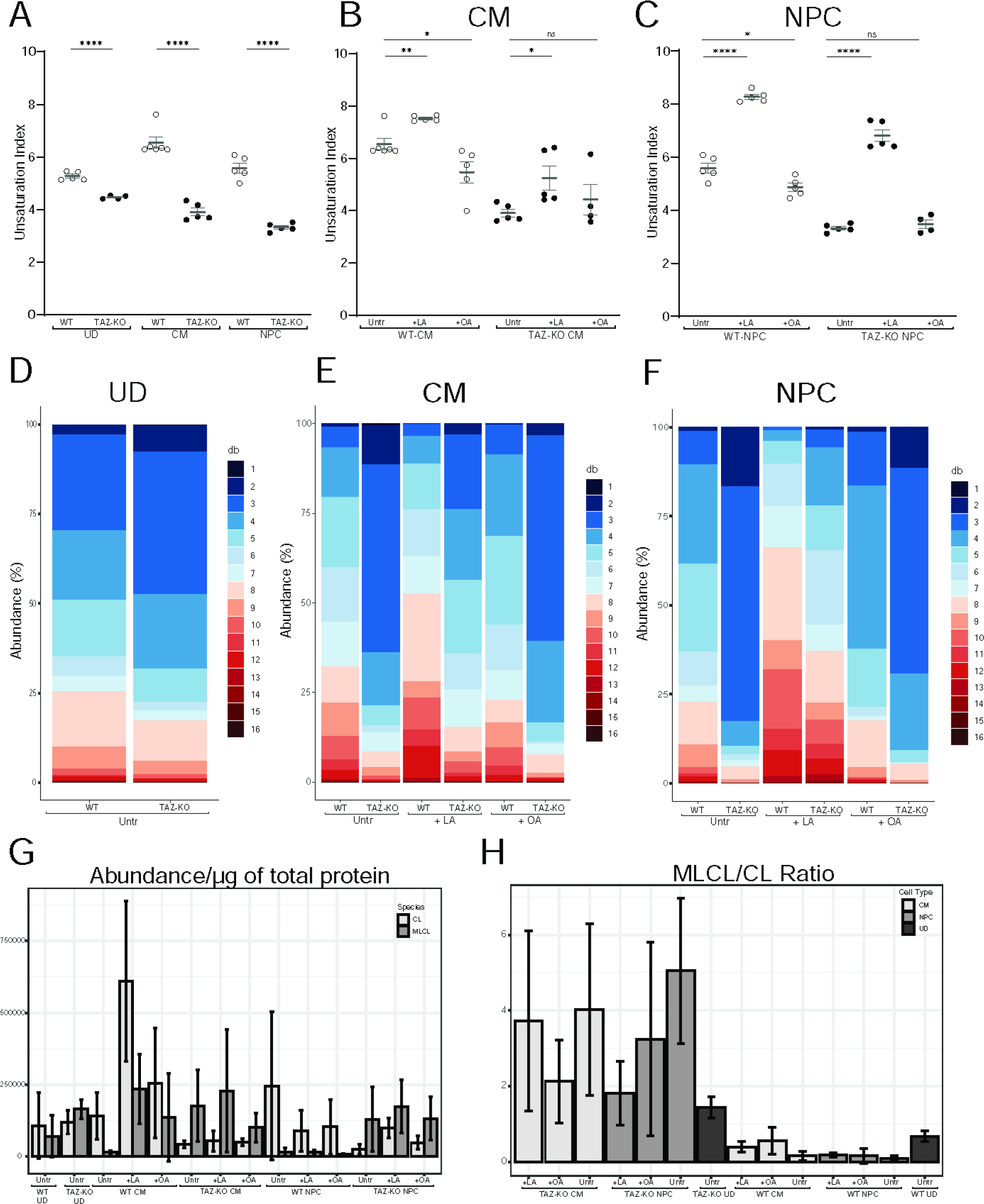
***CL acyl chain manipulation predictably alters CL saturation and species.*** (A) Baseline CL saturation index is reported in each cell type and genotype. (B) CL saturation index is reported in WT and TAZ-KO CMs treated with 100µM of linoleic acid (LA) or Oleic Acid (OA) for 8 days. (C) CL saturation index is reported in WT and TAZ-KO NPCs treated with 100µM of linoleic acid (LA) or Oleic Acid (OA) for 8 days. (D-F) The distribution of CL species containing specified numbers of double bonds was reported as percentage of total CL in untreated cells, and those supplemented with 100µM of LA or OA. (G) CL and MLCL abundance are reported per microgram of total protein. (H) MLCL/CL ratio of indicated cell types and genotypes. Significant differences are indicated; *≤0.05, **≤0.005, ***≤0.0005, and ****≤0.00005.

CL species fully saturated with LA or OA species would have 8 and 4 double bonds respectively. As expected, CMs and NPCs supplemented with LA showed increased unsaturation index (Fig. 5 B-C), as well as increased species with >3 double bonds (Fig. 5 E-F), which was less pronounced in TAZ-KO cell types compared to WT. Conversely, OA significantly lowered unsaturation index in WT CMs and NPCs (Fig. 5 B-C), while increasing CL species with fewer double bonds (Fig. 5 E-F).

### Shifting tissue-specific CL acyl content changes mitochondrial phenotypes

To address the impact on CL saturation on downstream phenotypes, we analyzed mitochondrial morphology phenotypes post FA supplementation in WT and TAZ-KO CMs and NPCs. FA supplementation had minimal impact on TAZ-KO CM mitochondrial morphology parameters, while WT CMs supplemented with LA or OA showed significant reductions in mitochondrial count, area, and branching (Fig. S8 A and C). Similarly, TAZ-KO NPCs showed minimal changes in response to CL saturation changes. However, WT NPCs had significant alterations in mitochondrial morphology, showing an inverse relationship between the effects of LA and OA, with LA causing significant increases in mitochondrial fragmentation and OA causing significant increases in mitochondrial network interconnectivity (Fig. S8 B and D).

Alterations in mitochondrial network connectivity can be indicative of changes in mitophagy, wherein mitochondrial fusion protects against, and mitochondrial fragmentation enables, mitochondrial engulfment by autophagosomes (68–69). However, changes in mitochondrial morphology after lipid supplementation were not associated with decreased mitochondrial mass as assessed by immunoblot analysis of TOM20 content (Fig. S8 E and F). While these data indicate that FA supplementation did not trigger mitophagy responses, analysis of the relative abundance of the active, lipidated form of autophagy protein LC3 revealed differences in overall cellular autophagy activity. Notably, FA supplementation resulted in increased LC3 activation in WT CMs and NPCs, but not TAZ-deficient CMs or NPCs (Fig. S9 A-B, D-E).

Taken together, these data indicate that, while WT CMs undergo significant remodeling of mitochondrial networks and show signs of overall autophagy activation in response to changes in lipid environment, TAZ-KO CMs are unable to do so.

### Characterizing OXPHOS complex assembly post CL acyl chain supplementation

To evaluate the effects of LA or OA supplementation on OXPHOS assembly, we performed blue native PAGE and immunoblotted for complexes I, II, III, and CIV alone or in combination (Total OXPHOS). We observed no significant alterations in CM or NPC complex assembly or abundance post treatment with either LA or OA (Fig. 10 and Fig. S11). Consistent with previous findings, TAZ-KO NPCs had reduced CIV present in the CIII2&CIV2 intermediate complex at baseline, which was partially restored after treatment with LA or OA (Fig S10).

## Discussion

We developed and characterized a TAFAZZIN-deficient iPSC model to study tissue-specific mitochondrial phenotypes in BTHS. In doing so, we expand the understanding of OXPHOS defects caused by TAZ-deficiency and highlight an emerging role for abnormal mitophagy in the cardiac pathology of BTHS.

In TAZ-KO CMs, we observed evidence of dysregulated mitophagy, including dysmorphic mitochondria/mitochondrial cristae, differential expression of key autophagy-associated genes, and an inability of TAZ-deficient CMs to properly initiate stress-induced mitophagy. Interestingly, we also observed the presence of PDH positive vesicles in TAZ-deficient CMs possibly representing upregulation of a compensatory, alternative mechanism of mitochondrial quality control mediated by mitochondrial-derived vesicles (MDVs) (62).

Our findings corroborate emerging evidence from BTHS cell and animal models that have recently suggested a role for mitophagy in the cardiac pathogenesis of BTHS (35). Prior studies demonstrated that TAZ-deficient mouse embryonic fibroblasts displayed impaired formation of mitochondria-engulfing autophagosomes (42). Intriguingly, recent studies in different *TAFAZZIN*-deficient mouse strains suggested that mitophagy capacity may be a strong modifier of cardiac pathology and outcome (35).

We hypothesize that defective mitophagy could provide the missing link to unify the prenatal and postnatal cardiac complications in BTHS, as it is both a critical component of CM differentiation and maturation as well as function (36, 43–44). We also saw evidence for abnormal mitophagy in TAZ-deficient NPCs, but these appeared to be less severe than what was observed in TAZ-deficient CMs. Although CMs and neuronal cells are both active in basal mitophagy, a diverse repertoire of regulatory mechanisms are responsible for maintaining homeostasis in each tissue type by carrying out mitochondrial remodeling during development and differentiation, post-stress induction, and other cell specific contexts (36). It is tempting to speculate that the diversity of mitophagy pathways, and the complex interplay of cellular and environmental cues, plays an important role in defining the tissue types affected in BTHS.

OXPHOS defects, including a reduction in supercomplex stability, have been implicated in several TAZ-deficient models (17, 28–30). We observed cell type-specific differences in OXPHOS assembly and activity, with TAZ-KO CMs having the greatest reduction in basal and maximal respiratory capacity, as well as ATP production and spare respiratory capacity. As mitochondrial complex assembly and expression are regulated in a tissue-specific manner and regulation is governed, in part, by reliance on OXPHOS vs glycolytic activity, continued investigations into differential fuel utilization, substrate flexibility, and respiratory capacity in mature TAZ-KO CMs and neurons will provide additional insight (49,51).

Notably, we found a significant reduction in CIV abundance and activity in the CIII-CIV tetramer specific to TAZ-KO NPCs. CIII-CIV intermediates have been shown to be preferentially stabilized in cell types favoring glycolysis, including in NPCs, and this interaction is known to be stabilized by CL (51–52). In fact, loss of CL results in the specific destabilization and loss of the CIV portion of the CIII-CIV tetramer (51–52). Additionally, CIII and CIV enzymatic activity has been observed to be increased nearly 2-fold as a member of the CIII-CIV complex rather than as free units, which may fine tune the efficiency of the electron transport chain (53–54). How the specific loss of CIV in the CIII-CIV intermediate complexes contributes to subtle changes in respiratory capacity and flexibility in TAZ-KO NPCs and neurons warrants further investigation.

CL tissue-specific acyl content adds an additional layer of complexity to these investigations, and recent evidence suggests tissue-available phospholipid pools are the defining factor governing CL acyl content, particularly driven by the opposing relationship between LA (18:2) and OA (18:1) side chains (13). Here, we show that CL acyl content can be manipulated *in vitro*, which predictably shifts CL acyl chain complement and saturation. Further, we found that shifts in CL saturation can have a dramatic impact on mitochondrial network morphology and mitophagy pathways in WT cells which is not recapitulated in TAZ-KO cell types. Importantly, post LA or OA treatment, CL levels remain significantly lower in TAZ-deficient cells, which results in less overall remodeled species compared to WT counterparts.

There remain many unanswered questions related to BTHS pathology including the implications of acyl content on the functions and binding interactions of CL, how TAFAZZIN, or other yet unidentified tissue-specific remodeling enzymes, remodel CL from the available PL pool, how the CL complement in each tissue supports both normal function and drives downstream dysfunction, and how MLCL functions in these pathways in the absence of CL. Although tissue-specific CL content represents an enticing candidate for therapeutic manipulation in BTHS, studies in mice have shown that strains with variable BTHS phenotype have similar alterations in CL and MLCL content, and genetic modifiers likely act downstream of CL abnormalities (35).

Limitations of this study include that investigations utilizing directed differentiation of iPSCs grown in monolayer culture do not mimic the complex cellular organization of mature tissues nor the interplay of tissues and the environment. However, this work can serve as a reference for further investigations in iPSC derived mature cardiomyocytes and neurons as well as tissues derived from TAZ-deficient murine models (35).

Taken together, our results have implications for understanding the tissue-specific pathology of BTHS and potential for tissue-specific therapeutic targeting. Moreover, our results highlight an emerging role for mitophagy in the cardiac pathophysiology of BTHS and reveal a potential neuron-specific bioenergetic phenotype.

## Materials and Methods

### Cell lines and cell culture conditions

iPSC WT cells were purchased from Coriell Institute (GM26105). Cells were grown at standard culturing conditions of 37°C, 5% CO2. iPSC WT and TAZ-KO undifferentiated cells were maintained in Essential 8 medium (Thermofisher A1517001) on plates coated with Geltrex reduced growth factor basement membrane (Thermofisher A1413302). For routine passaging, cells were detached with StemPro Accutase (Thermofisher A1110501) and replated in E8 containing ROCK inhibitor (Y-27632) (BD Biosciences 562822). After 24 hours, media containing ROCK inhibitor was replaced by E8 for the remainder of experiment. Mycoplasma contamination was routinely monitored and not detected.

### CRISPR/Cas9 genome editing

sgRNAs targeting exon 2 of the *Tafazzin (TAZ)* gene were designed at www.crispr.mit.edu (Supplemental Fig S1). Guides were selected based on the scoring algorithm detail in Hsu et al. 2013 (55). Synthesized sgRNA_H1 and sgRNA_R1 were individually cloned into pSpCas9(BB)-2A-Puro (PX459) V2.0 vector. pSpCas9(BB)-2A-Puro (PX459) V2.0 was a gift from Feng Zhang (Addgene plasmid # 62988 ; http://n2t.net/addgene:62988 ; RRID:Addgene_62988 (56). WT iPSC cells were transfected with both guide RNA vectors using Lipofectamine 2000 (Thermofisher 11668019). 24-hours after vector transfection, cells were subjected to Puromycin selection (2 µg/mL) for 48 hours. Following puromycin selection, cells were passaged to isolate single cell clones. Confluent colonies derived from single cell clones were collected using a cloning cylinder and expanded for DNA isolation and screening. The region surrounding the CRISPR/Cas9 target site was PCR amplified using AccuPrime GC-Rich DNA Polymerase (Thermofisher 12337024) (Forward Primer: 5’ TACATGAACCACCTGACCGT 3’, Reverse Primer: 5’ CTGAAACTCCGCCACATCTGG 3’). PCR products were resolved on a 3% agarose gel. Four TAZ-KO clone isolates, c3, c10, c13, and c24, were generated with a 50 bp deletion (c.456_505del, p.Lys56_Met73). For these 4 TAZ-KO clones isolates, we amplified and Sanger sequenced 2 of the top 5 predicted off-target sites per sgRNA, which revealed no detectable off-target CRISPR/Cas9 genome editing activity. None of the top 10 predicted off-target sites were in coding regions. After testing differentiation efficacy, we moved forward with iPSC TAZ-KO clone 13 for future experiments.

### iPSC differentiation into cardiomyocytes

iPSC WT and TAZ-KO cardiomyocytes (CMs) were differentiated as described by Cho et al. (57). Briefly, iPSCs were seeded on Geltrex in 6-well tissue culture plates in E8 medium. At confluency, marking day 0 of CM differentiation, cells were treated with 4 µM CHIR-99021 (Cell Signaling Technologies) in RPMI Glutamax (Thermofisher 61870127) supplemented with B27 Supplement minus insulin (B27-) (Thermofisher A1895601) for 48 hours. On day 2, cells were treated with 2 µM CHIR-99021 in RPMI- B27- for 24 hours, followed by 48 hours of 5 µM IWR-1-endo (Stem Cell Technologies 72562) on day 4. On day 6, cells were placed in B27- for 48 hours and subsequently cultured in RPMI Glutamax supplemented with B27 Supplement without vitamin A (B27+) (Thermofisher 12587010) until collected. After beating was achieved, cells were transiently cultured in a selection media containing no glucose, supplemented with 4mM sodium lactate (Sigma L7022) for 48 hours.

### iPSC differentiation into neural progenitor cells

WT and TAZ-KO iPSCs were differentiated into neural progenitor cells (NPCs) using PSC Neural Induction Medium Kit (Thermofisher A1647801) as described by the manufacturers protocol. NPCs were passaged and used at P1-P5 for experiments.

### Lipid treatment

CMs, and NPCs were grown in standard culturing media, B27+ and PSC Neural Expansion media, respectively, until desired confluency was reached. Respective medias were made containing either 100µM or 250µM Linoleic acid albumin (Millipore Sigma L9530) or Oleic Acid albumin (Millipore Sigma O3008). Cells were treated with respective lipid-containing medium every 48 hours for 8 days, after which they were collected for corresponding analyses.

For lipid treated cell survival cells were plated in 96 well plates and undertook a course of either no treatment, 100µM or 250µM LA or OA for 8 days. Cells were counted via the Celigo Cytometer (Nexcelom Biosciences) at days 0, 2, 4, 6, and 8. Each graphed point represents >3 wells analyzed per day and aggregated.

### RT-PCR and quantitative RT-PCR

For RNA extraction, a cell pellet >3x10^6^, or 1-well of a 6-well plate, was resuspended in 500µL of TRIZOL (Thermofisher 15596018). RNA was extracted from cells in TRIZOL using the RNeasy Plus Kit (Qiagen 74134) according to the manufacturer’s instructions. cDNA was synthesized from extracted RNA using the iScript cDNA Synthesis Kit according to the manufacturer’s suggested protocol, using 1µg of RNA (BioRad 1708891). cDNA was further diluted 1:10 in RNase free diH20. PCR validation of cell type specific markers was performed for UDs iPSCs, iPSC-CMs, and iPSC-NPCs using primers for corresponding marker genes. PCRs were run under standard conditions and products were resolved on a 2% agarose gel. For quantitative RT-PCR, 12µL reactions were generated using the SsoAdvanced Universal SYBR Green Supermix (Bio-Rad 1725271) as per the manufacturer’s instructions and included: 2.4µL of cDNA as well as each respective forward and reverse gene-specific primers. Each sample-primer reaction pair was plated in triplicate per run. Each plate included two internal controls: both a no reverse-transcriptase control (No-RT) for each cDNA sample, and no template controls (No-Template) for each primer pair. Reaction conditions for qRT-PCR were as follows: 2 min at 95°C, followed by 40 two-temperature cycles of 5 s at 95°C and 30 s at 60°C.

### Immunofluorescence

For immunofluorescence, cells were plated onto Geltrex coated 18 mm glass coverslips (VWR, cat.no. 48380046) or ibidi chamber slides (8 chamber or 12 chamber). At the indicated timepoints, cells were fixed in 4% paraformaldehyde (EMS 157-4) for 30 minutes. Cells were then permeabilized in 0.5% Triton-X100 for 5 minutes, washed, and blocked in 2.5% donkey serum for 1 hour. Following, coverslips were incubated in 2.5% donkey serum containing primary antibody for 3 hours. Subsequently, coverslips were incubated in 2.5% donkey serum containing secondary antibody for 30 minutes. After final washing steps, nuclei were stained with DAPI solution (Invitrogen, cat.no. R37606) for 10 minutes and then coverslips mounted onto glass slides with ProLong Gold Antifade Mountant (Invitrogen P36930). Mounted coverslips were allowed to dry for at least 24 hours prior to imaging. For chamber slides, chambers were removed, and a glass coverslip was mounted using Prolong Gold reagent. Cells were imaged using a Leica SP8 confocal microscope and a Leica THUNDER widefield deconvolution fluorescence microscope, equipped with a Leica K8 Scientific CMOS camera, an LED8 solid-state light source, and a 63X (1.4 N.A.) oil immersion objective.

### Live cell mitophagy assay

For live cell fluorescence mitophagy assays, the Dojindo Molecular Technologies Inc Mitophagy Detection Kit (Dojindo Molecular Technologies Inc MD0110) was used as per the manufacturer’s protocols. In brief, cells were co-stained with the Mtphagy Dye and the Lyso Dye and imaged using a Leica SP8 confocal microscope. Colocalization of Mtphagy and Lyso Dyes was analysed in Fiji (70) using the Coloc 2 plugin, and normalized to cell number. For untreated WT and TAZ-KO CMs, 94 and 88 cells were analyzed respectively. For WT and TAZ-KO CCCP treated CMs, 75 and 88 cells were analyzed, respectively. For WT NPCs 194 cells were analyzed in the untreated group, and 146 in the CCCP treated group. For TAZ-KO NPCs 277 and 209 cells were analyzed in the untreated and CCCP treated groups, respectively.

### Mitochondrial morphology

Cells of indicated type and genotype were subjected to immunofluorescence staining of TOM20 and imaged using a DeltaVision Elite high-resolution fluorescence deconvolution microscope, equipped with a Scientific CMOS camera, an UltraFast solid-state illumination, and a 60X (N.A. 1.42) oil immersion objective. Images were analyzed using the Fiji plugin ‘Mitochondria Analyzer’ version 2.1.0 (48) following steps outlined in the ‘Mitochondria Analyzer’ user manual. In brief, first adaptive thresholding was utilized determine optimal thresholding parameters to faithfully capture mitochondrial morphology, discrete mitochondria, and eliminate background. Distinct optimal thresholds were chosen for each cell type, i.e., iPSC-NPCs vs iPSC-CMs. Following thresholding, images were analyzed for all parameters using the ‘Analysis’ command on a per-cell basis. Descriptors of each individual parameter can be found in the user manual. For each condition, one point plotted represents an individual cell’s measurements.

### RNA Sequencing

RNA-seq was performed on WT and TAZ-KO UD iPSCs in triplicate on three biological replicates that represent independent collections and differentiations. RNA was isolated as described above for all samples on the same date and time to mitigate batch effects. RNA concentration and purity were measured via the NanoDrop One (Thermofisher) prior to sample submission. Whole transcriptome library preparation, Illumina sequencing (Illumina HiSeq), and analyses were performed by Novogene Bioinformatics Technology Co., LTD. In brief, reads were generated and mapped to Hg38 using HISAT2 and reads aligning to each gene were identified via Feature Counts. Reads were trimmed to remove linker sequences, and low-quality reads with uncertain nucleotides constituting >10% (N>10%) or low-quality nucleotides (base quality less than 5) constituting >50% were removed. Fragments per kilobase of transcript per million fragments mapped (FPKM) of each gene were calculated and differential expression analysis between cell types was performed with DESeq2 R package described in Anders et al., 2010 (71). Genes with an adjusted p-value < 0.05 and a |log2(FoldChange)| > 0 were considered differentially expressed between genotypes. Enrichment analysis was performed on differentially expressed genes using ClusterProfiler to identify KEGG pathways and Gene Ontology (GO) Terms.

### Whole Cell Lysis Extraction

An aliquot of >3x10^6^ cells, or 1-well of a 6-well plate, was collected and centrifuged for 4 min at 1000 rpm. The resulting pellet was washed twice with ice-cold PBS and re-centrifuged to pellet cells. Cells were lysed with RIPA lysis buffer (1% (v/v) Triton X-100, 20 mm HEPES–KOH, pH 7.4, 50 mm NaCl, 1 mm EDTA, 2.5 mm MgCl2, 0.1% (w/v) SDS) spiked with 1 mm PMSF for 1 hour at 4°C with rotation. Insoluble material was removed by centrifugation for 30 min at 21,000g at 4°C. The supernatant was collected, and protein content was quantified using a bichichronic acid (BCA) assay (Pierce 23225). Aliquots were made for immunoblotting and stored at -80°C.

### Mitochondrial Isolation

Isolation of mitochondria from iPSC cell types was performed as previously described by Lu et al., and Jha et al., (9, 58).

### Immunoblotting

Aliquoted micrograms of whole cell extracts or mitochondria were resuspended in XT Sample Buffer (BioRad 1610791) and Reducing Agent (BioRad 1610792). Proteins were resolved on Criterion XT 12% Bis-Tris gels (BioRad 3450117) in XT MOPS Running buffer (BioRad 1610788). Proteins were blotted to Immuno-Blot PVDF (BioRad1620177) membranes using a liquid transfer system at 100 V for 1 hour. Following transfer, PVDF membranes were blocked with 5% (w/v) milk, 0.05% (v/v) Tween-20 in PBS for 1 hour at room temperature or at 4°C if longer, with rocking. Membranes were briefly washed with PBST (PBS with 0.2% (v/v) Tween-20) and incubated with primary antibody in PBST with 0.02% (w/v) Na-Azide overnight at 4°C with rocking. Following three 10 minute washes with PBST at room temperature, HRP-conjugated secondary antibodies were added and incubated for 1 hour at room temperature. Following three washes for 10 minutes with PBST and two washes for 10 minutes with PBS, immunoreactive bands were visualized using SuperSignal West Pico PLUS (Thermofisher 34578). For chromogenic/colorimetric blots the WesternBreeze Chromogenic Kit, anti-rabbit or anti-mouse kit (Invitrogen) was used, which contains blocking solutions, wash buffers, secondary antibodies, and a chromogenic solution. Images were captured using a ChemiDocMP (Biorad) Band quantification was performed using ImageJ and protein expression values were normalized to loading controls.

### Antibodies

Antibodies against the following proteins were used for immunoblotting: β-actin (Life Technologies M4302), TAZ (#2C2C9, Epitope: 237-TDFIQEEFQHL, exon 11) (9), LC3 A/B antibody (Cell Signaling Technologies 4108), SDHA (Abcam 14715), and GRP75 (Antibodies Inc., 75-127). Two HRP-conjugated secondary antibodies were used: goat anti-rabbit (Abcam ab6721), goat anti-mouse (Abcam 205719). The following antibodies were used for immunofluorescence staining of cell type-specific markers in cardiomyocyte and neural progenitor cells, respectively: Troponin T (Epredia MS- 295-P1) and Nestin (Millipore MAB5326). An antibody against TOM20 (Santa Cruz Biotechnology sc- 17764), was used for immunofluorescence staining of mitochondria. Antibodies against the following proteins were used for Blue Native Page: NDUFB8 (CI, Thermofisher 459210), SDHB (CII, Thermofisher 459230), MTCO1 (CIV Thermofisher 459600), and OxPhos Human WB antibody cocktail (Thermofisher 458199). For visualization, Western Breeze Chromogenic detection kit (α-mouse, Thermofisher WB7103) was used.

### 1D Blue Native Page

One dimensional blue native page was carried out as previously described in (9, 58–59). Briefly, 1x10^6^ cell pellets were solubilized for 20 minutes on ice in sample buffer cocktail containing native page sample buffer (Thermofisher BN2003) with a digitonin-protein ratio of 8g/g. Extracts were clarified by a 10 minute centrifugation at 20,000*xg* at 4°C and analyzed by 1D BN/SDS PAGE.

### In gel activity assay

Clear native page followed by complex I and complex IV in gel activity were assessed as previously described by Jha et al. and Milenkovic et al. (58, 60).

### Lipidomics Dataset 1

Lipids were extracted from cell pellets (3x10^6^) of WT and TAZ-KO UD-iPSCs and analyzed as previously described in Anzmann et al. (17). See data in Fig. 1.

### Lipidomics Dataset 2

Cell pellets of 1x10^6^ lipid supplemented WT and TAZ-KO UD, CM, and NPCs, and untreated controls, were analyzed as described in Oemer et al. and Wohlfarter et al. (13, 60). See data in Fig. 4 and supgplemental material.

### OXPHOS function

Cells (UD iPSCs and NPCs) were plated in geltrex coated 96-well Seahorse plates (Agilent 103794- 100) and allowed to grow until confluency was reached, iPSC-CMs were differentiated from UDs in geltrex coated seahorse plates until beating was achieved. The Agilent XF Mito Stress Test kit (Agilent 103015-100) was used to measure oxygen consumption and extracellular acidification on the Seahorse XFPro 96well Analyzer (Agilent). Cells were incubated in assay medium containing NucBlue Live Cell Stain ReadyProbes (Invitrogen R37605) and were counted prior to the Seahorse assay using a Celigo Cytometer (Nexcelom Biosciences). Assay data (OCAR and ECAR) was normalized per 1000 cells using the Seahorse Wave Desktop Software.

### Electron Microscopy

Transmission electron microscopy (TEM) and immunoelectron microscopy (IEM) were performed at the Johns Hopkins Microscopy Core. For TEM cells were fixed (in plates) in 2% paraformaldehyde and 1% glutaraldehyde (Electron Microscopy Sciences) in 0.1 M sodium cacodylate buffer (pH 7.4) for 1 hour at room temperature, followed by three washes in 0.1 M cacodylate buffer. Then, cells were post-fixed for 1 hour in 1% osmium tetroxide (Electron Microscopy Sciences) in the same buffer at room temperature. Samples were washed in water and stained for 1 hour at room temperature in 2% uranyl acetate (Electron Microscopy Sciences), washed again in water and dehydrated in a graded series of ethanol. Then, samples were embedded in Embed-812 epoxy resin (Electron Microscopy Sciences). Ultrathin (50-60 nm) sections were cut using an Ultracut ultramicrotome (Reichart-Jung) and collected on formvar- and carbon-coated nickel grids, stained with 2% uranyl acetate and lead citrate. Samples were imaged on a Hitachi H7600 electron microscope under 80 kV. Immunogold samples were fixed in 1% glutaraldehyde (EM grade) 100 mM phosphate buffer 5 mM MgCl2 pH 7.4 for 1 hr and prepared as described above. Sections were stained with primary antibodies against TOM20 (Invitrogen MA34964) and PDH (Abcam 110333) and then labeled with secondary antibodies conjugated to 12nm and 6nm diameter gold particles respectively.

### Data Analysis

All experiments were performed in triplicate, with at least three biological replicates for each genotype and cell type. All graphical depictions and statistical analyses were carried out in GraphPad PRISM version 9.5.1. Between group comparisons were performed using unpaired T-test.

## Acknowledgments

We would like to thank Emmanouil Tampakakis and Vasiliki Machairaki for their generous guidance in iPSC cardiomyocyte and neuron differentiation respectively. We thank Nanami Senoo of the Claypool Laboratory for valuable discussion and technical support. We thankfully acknowledge use of instruments at the Light Microscopy Facility of the Department of Molecular Microbiology and Immunology at the Johns Hopkins Bloomberg School of Public Health. We thank the technical staff of the Johns Hopkins School of Medicine Microscopy Facility for their assistance with electron microscopy.

## Author contributions

Conceptualization, O.S.C., H.J.V., M.K., A.H.-B., and, S.M.C.; Methodology, O.S.C., K.W.H., Y.W., K.L., G.B., A.A., S.M.C., A. H-B., M.K., H.J.V.; Formal Analysis, O.S.C., K.W.H., Y.W., K.L., G.B., A.A., S.M.C., A. H-B., M.K., H.J.V. ; Investigation, H.J.V., A.H.-B., M.K., and, S.M.C.; Data Curation, O.S.C., K.W.H., Y.W., K.L., G.B., A.A.; Writing – Original Draft, O.S.C. and H.J.V; Writing – Review & Editing, O.S.C., K.W.H., Y.W., K.L., G.B., A.A., S.M.C., A. H-B., M.K., H.J.V. Supervision, H.J.V., A.H.-B., M.K., and, S.M.C.; Funding Acquisition, A.A., H.J.V.

## Funding and additional information

Research reported in this publication was supported by the National Heart, Lung, and Blood Institute of the National Institutes of Health under Award Number F31HL147454 to A. F. A as well as NIGMS (T32GM148383), Maryland TEDCO awards 2002-MSCRFD-5837 to H.V. and 2018-MSCRFD-4237 to H.V.

