## Supplemental table 1 for "Stem cell models of TAFAZZIN deficiency reveal novel tissue-specific pathologies in Barth Syndrome"

Supplemental Materials:

A

| sgRNA | Guide Sequence (5' to 3') | Off-Targets (5' to 3') | Score[1] | MMs[2] | Hg38 Location | Sequenced |
|---|---|---|---|---|---|---|
| | | TCAGAGTTCATTGAGAAGCGAAG | 0.5 | 4 | chr12:-6890445 | Y |
| | | CACAAGCTCACCAAGAAGCGTGG | 0.3 | 4 | chr5:+110321962 | Y |
| H1 | GAGATGAGGGTCGTCCATGCAGG | CACGAGCTCCTGGAGAAGCAAGG | 0.3 | 4 | chr19:-11912726 | N |
| | | TTCAGGCTCATCGAGGAGCGCAG | 0.3 | 4 | chr8:-104093813 | N |
| | | TATGAGCTCCTCAAGAAGCCCAG | 0.2 | 4 | chr12:-5342876 | N |
| | | GAGATGTAGCTCGTCCATGCTGG | 1.5 | 3 | chr3:+173436340 | Y |
| | | GTTCAGAGGGTCGTCCATGCAAG | 1.3 | 4 | chr7:-70923313 | Y |
| R1 | TACGAGCTCATCGAGAAGCGAGG | GATATGAGGGGAGTCCATGCAGG | 0.7 | 3 | chrX:-12667869 | N |
| | | GACAAGAGGTTGGTCCATGCCAG | 0.6 | 4 | chr7:+65692780 | N |
| | | CTGGTGAGGGTCTTCCATGCCAG | 0.6 | 4 | chr8:-125971306 | N |

[1] Off-target score calculated at cripsr.mit.edu based on scoring algorithm from Hsu et al. 2013
[2] The number of mismatches between the guide sequence and the "off-target" sequence
[3] PAM site is bolded

B

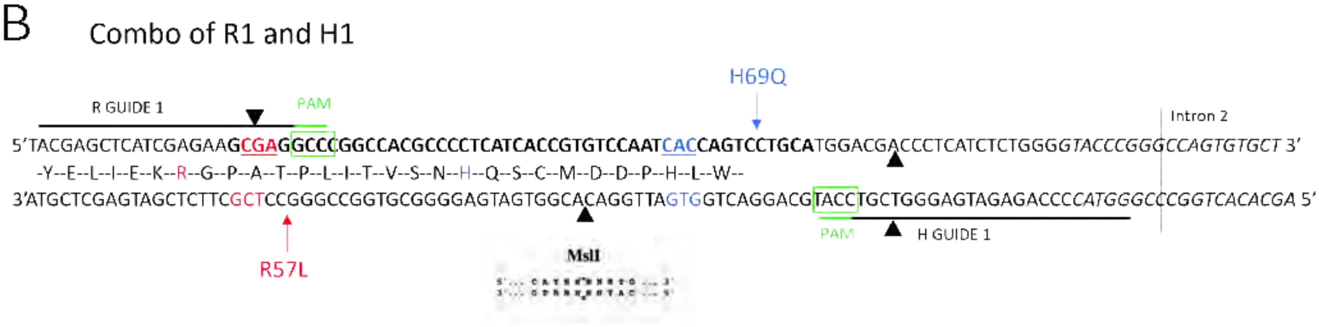


**Fig. S1.** ***Generation of a novel TAFAZZIN-deficient iPSC model***. (A) Two single guide RNAs used for the CRISPR TAZ-KO are reported, along with their predicted off target effects. (B) A representative schematic of the binding of the two single guide RNAs, near to two identified patient variants R57L and H69Q.

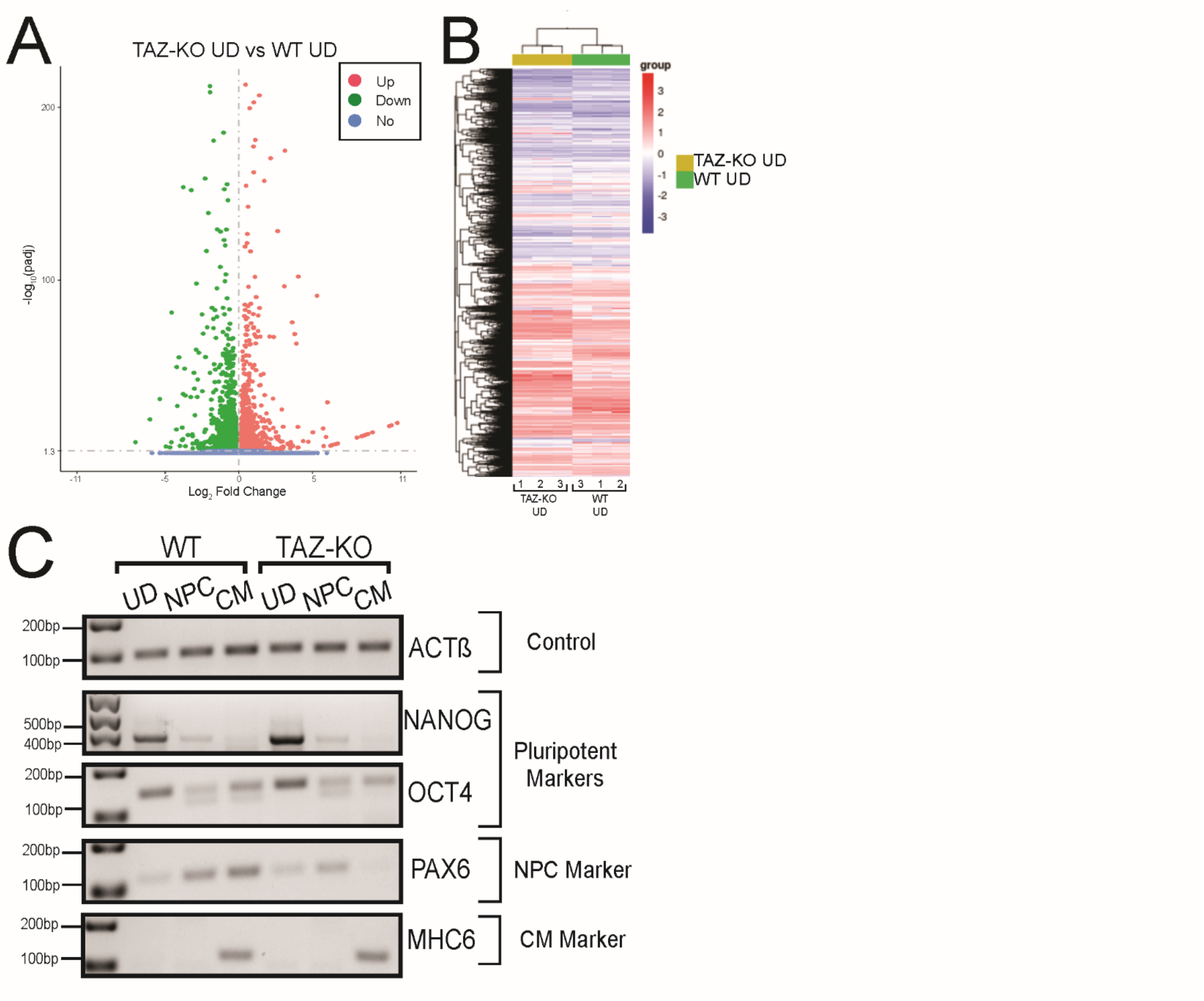


**Fig. S2.** ***RNAseq and validation of iPSC derived CMs and NPCs.*** (A) Volcano plot in UD WT and TAZ-KO cells demonstrate differentially expressed genes either upregulated (pink) or downregulated (green) in TAZ-KO genotype compared to WT. (B) Heatmap demonstrating expression and clustering of UD WT and TAZ-KO iPSCs. (C) RT-PCR for indicated cell type specific marker gene expression in CMs and NPCs.

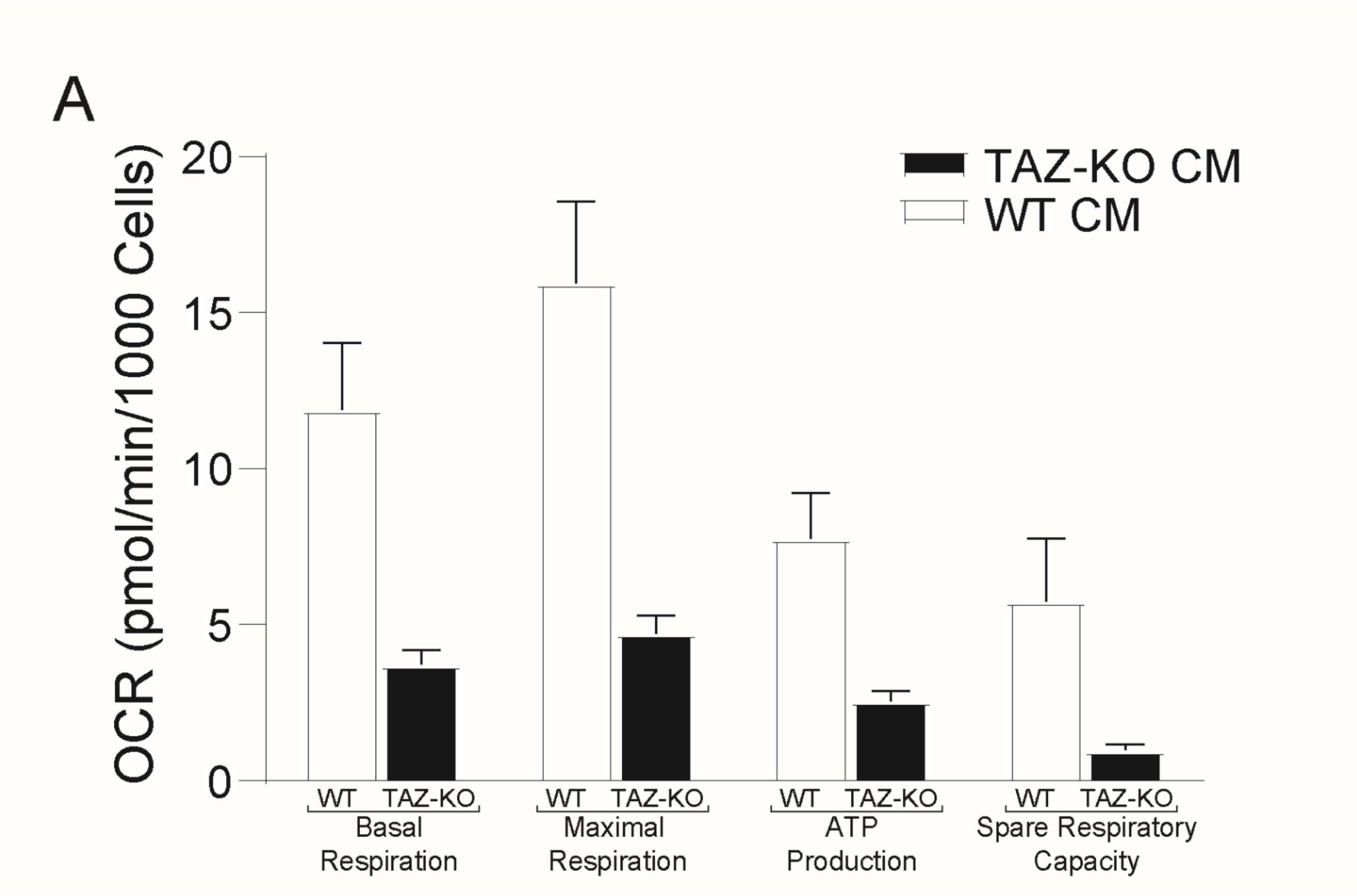


**Fig. S3.** ***TAZ-deficient CMs have reduced OXPHOS measures.*** (A) TAZ-KO CMs have significantly reduced basal respiration, ATP production, and spare respiratory capacity compared to WT counterparts.

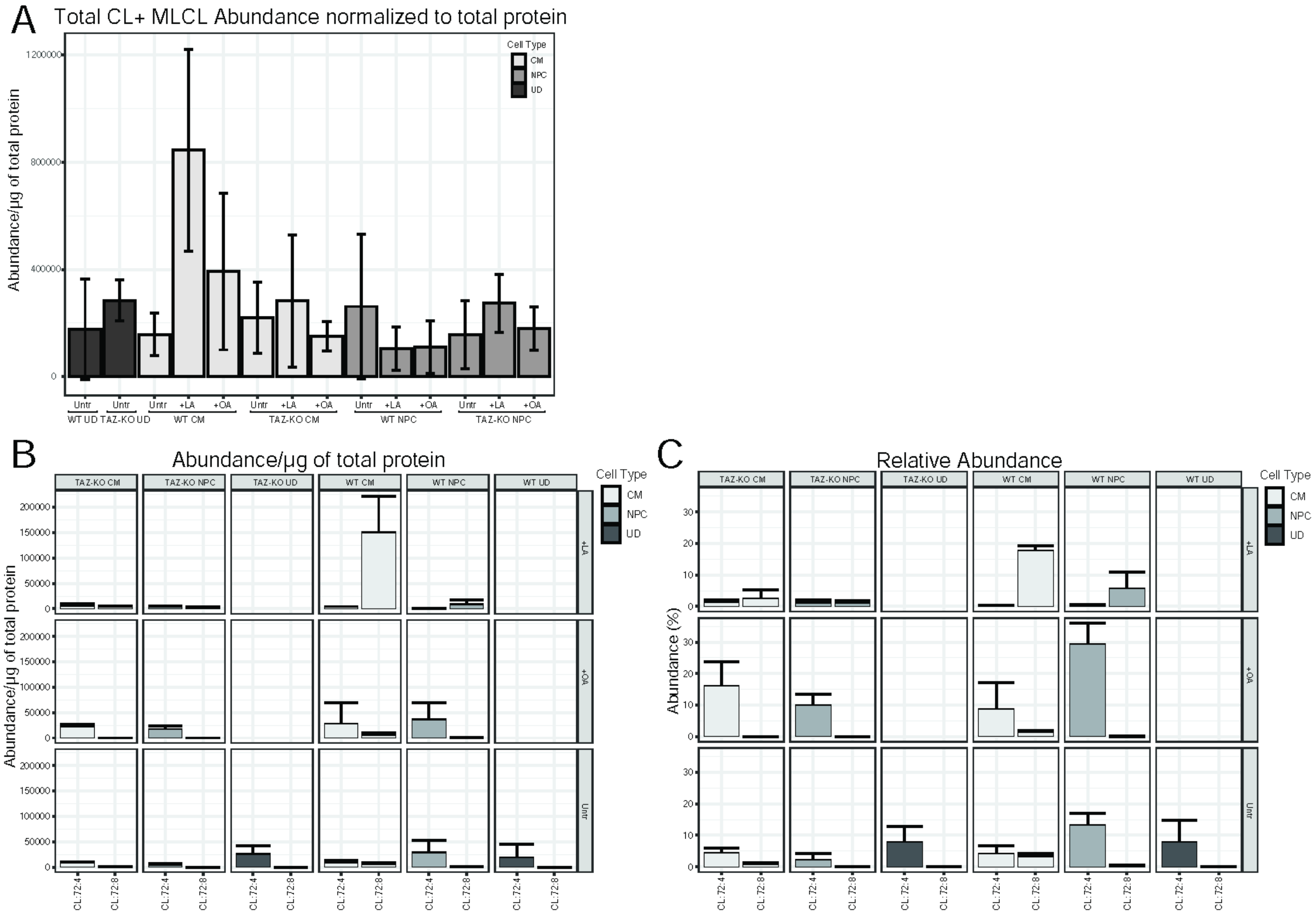


**Fig.S4.** ***Additive CL abundance and evaluation of CL(18:1)$_4$ and CL(18:2)$_4$ with lipid supplementation.*** (A) CL and MLCL abundance shown as a sum in each cell type-genotype pair with or without 100µM LA or OA supplementation. (B) Abundance and (C) Relative abundance of *CL(18:1)$_4$ and CL(18:2)$_4$* in untreated and 100µM lipid supplemented cells of each genotype.

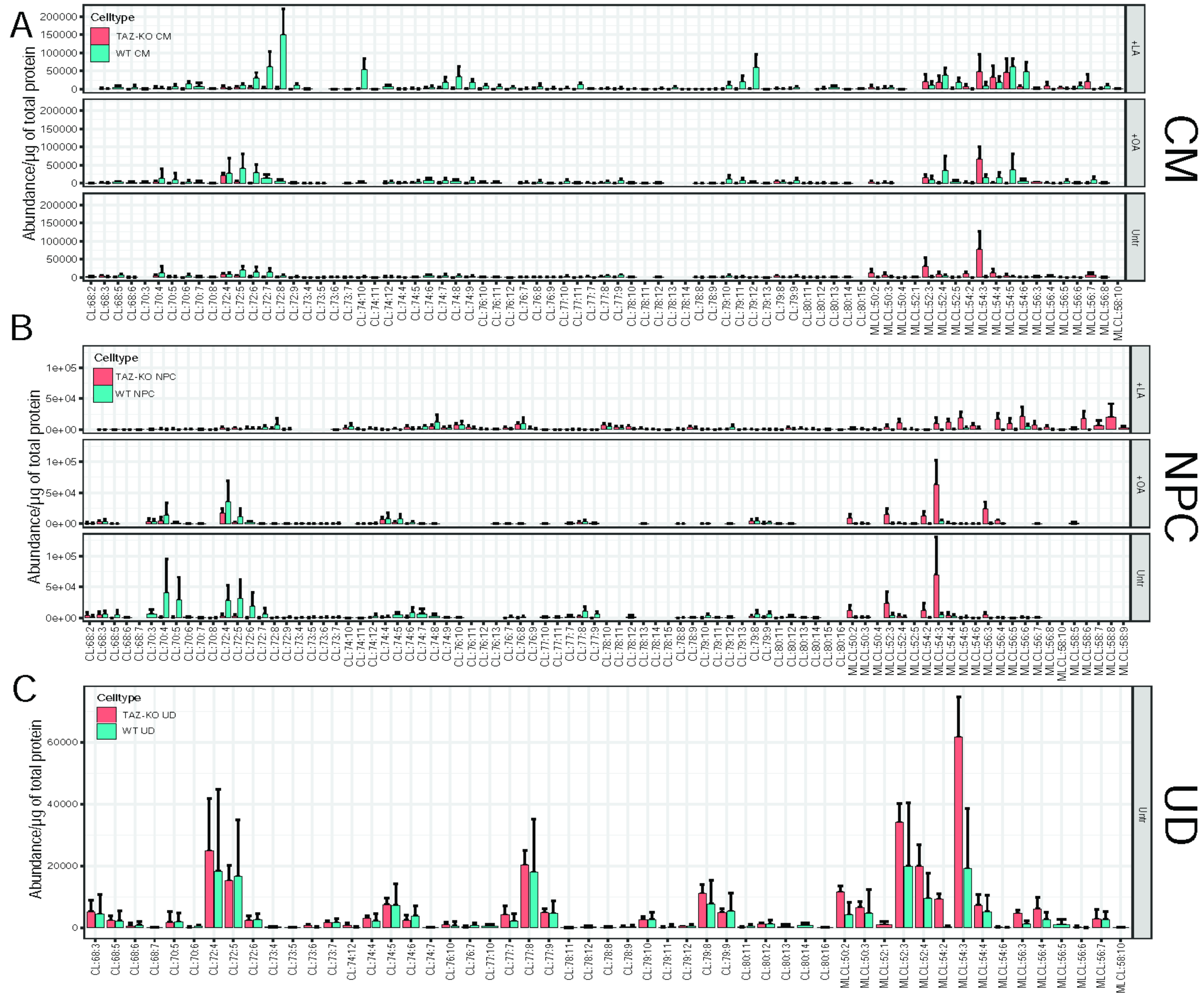


**Fig. S5.** ***CL species change with lipid supplementation.*** Distribution of all CL species measured for WT (blue) and TAZ-KO (pink) (A) CMs (B) NPCs, and (UDs). Treatment with 100μM LA or OA is indicated for CMs and NPCs.

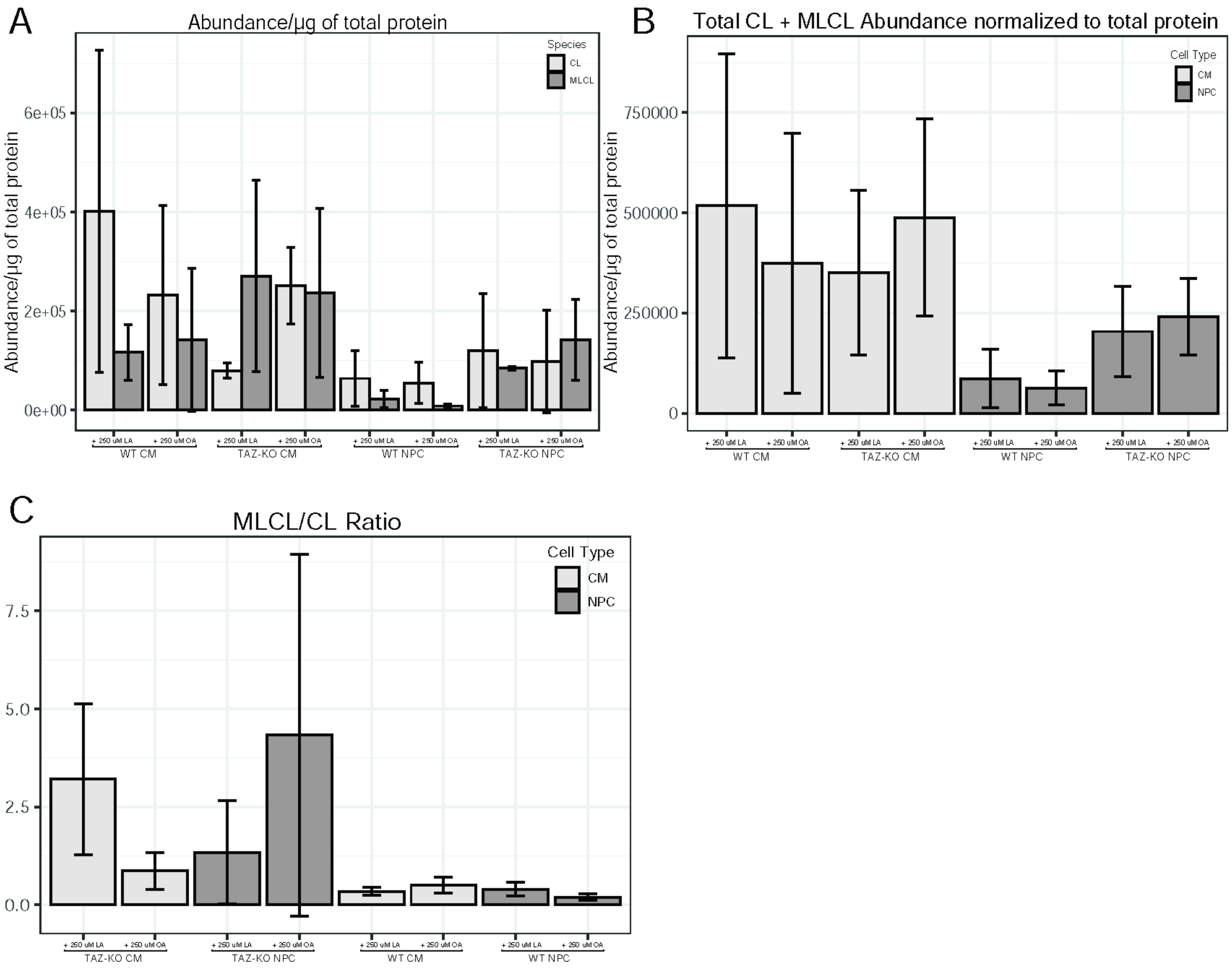


**Fig. S6.** ***250μM supplementation of LA or OA shows similar trends to 100 lipid μM.*** Cells were supplemented with either 250μM of LA or OA. (A) CL and MLCL abundance were measured and normalized to micrograms of total protein. (B) The additive MLCL and CL abundance are plotted normalized to total protein. (C) MLCL to CL ratio is reported for each cell type-genotype pair.

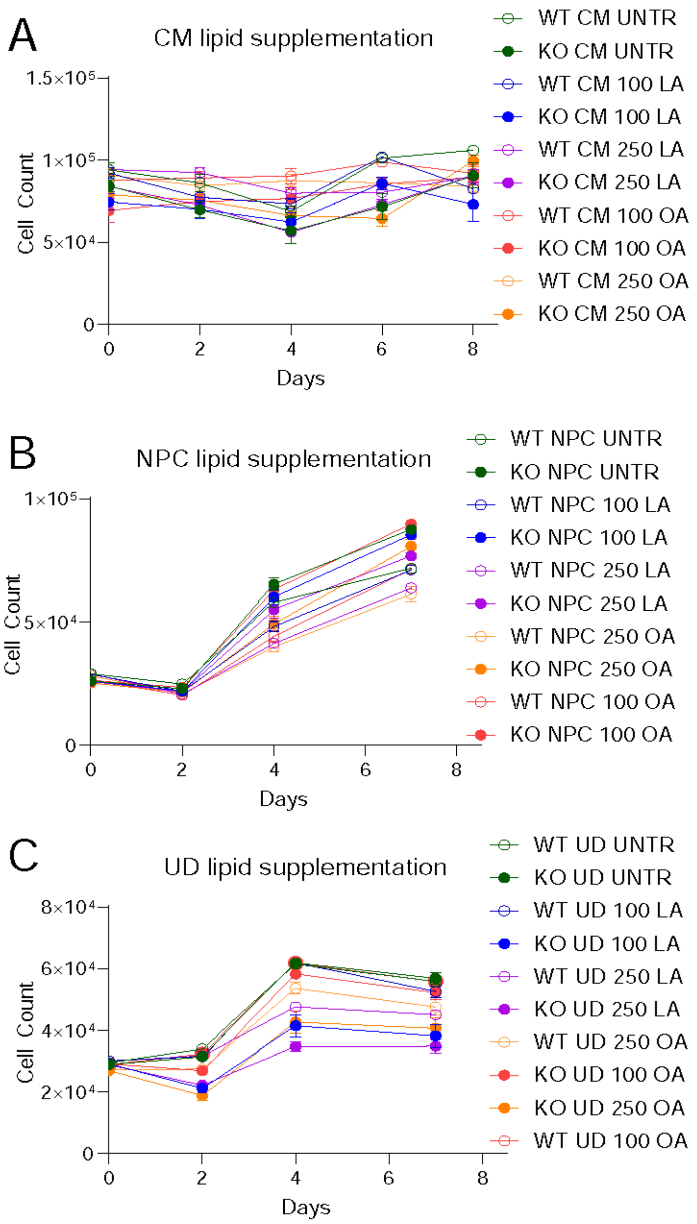


**Fig. S7.** ***Lipid supplementation does not alter cell growth and survival in CMs and NPCs.*** Each cell type-genotype pair were left untreated, or supplemented with 100µM LA or OA, or 250µM of LA or OA. Cell counts were measured in (A) CMs, (B) NPCs, and (C) UDs, every 2 days starting at day 0 and continued through day 8. Each count represents >3 technical replicates.

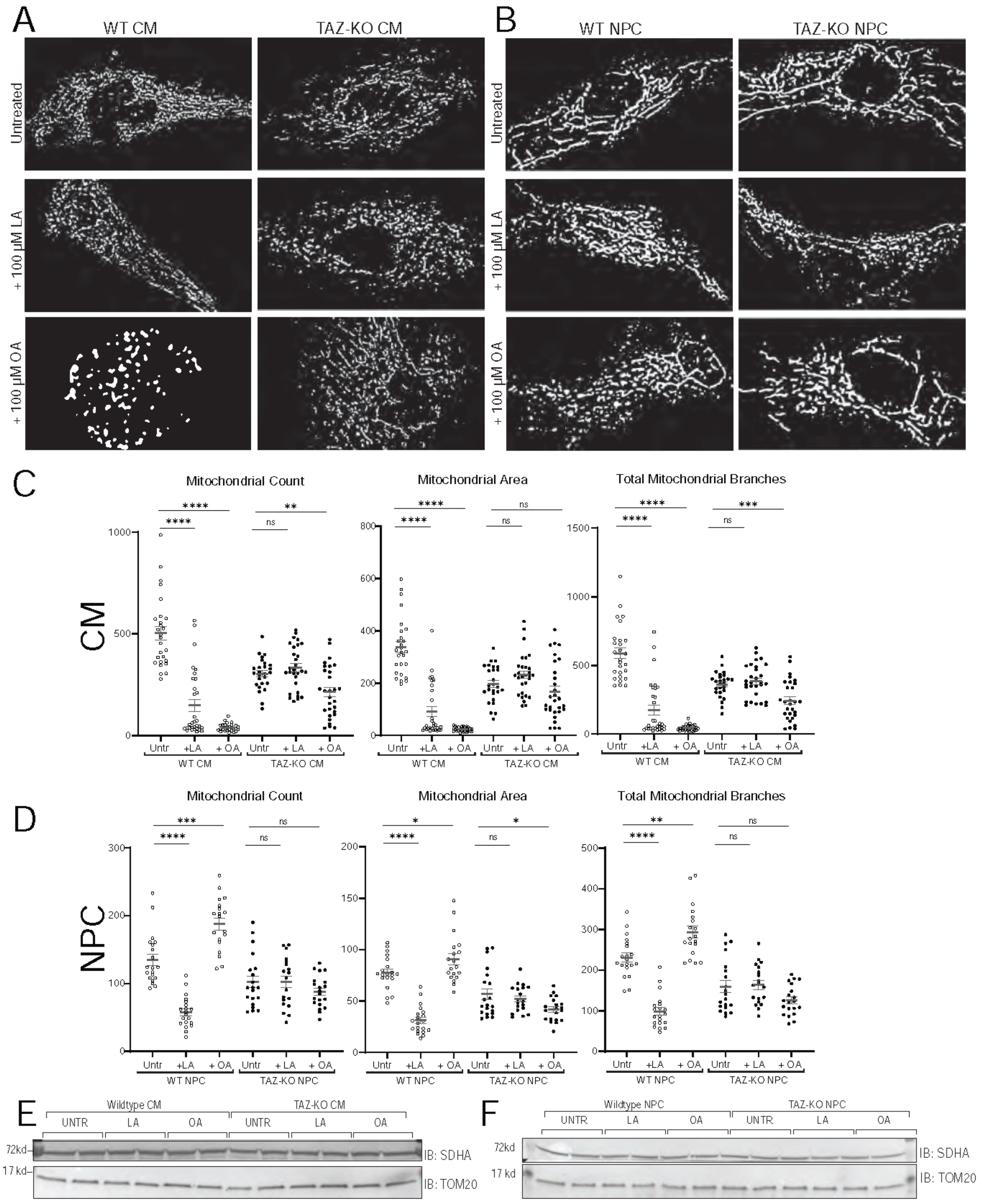


**Fig. S8*. Lipid treatment induces mitochondrial remodeling in WT but not TAZ-KO CMs and NPCs.*** (A-B) Representative immunofluorescence images of both untreated, and 100µM LA or OA supplemented WT and TAZ-KO (A) CM and (B) NPC. TOM20 stained mitochondria were segmented using Mitochondrial Analyzer. (C-D) Mitochondrial networks from individual cells were quantified using Mitochondrial Analyzer and metrics from individual cells are reported as well as the mean, and standard error of the mean. (E-F) Whole cell lysate (60 µg) of indicated cell lines and treatments were immunoblotted for TOM20 and SDHA. Significant differences are indicated; *≤0.05, **≤0.005, ***≤0.0005, and ****≤0.00005.

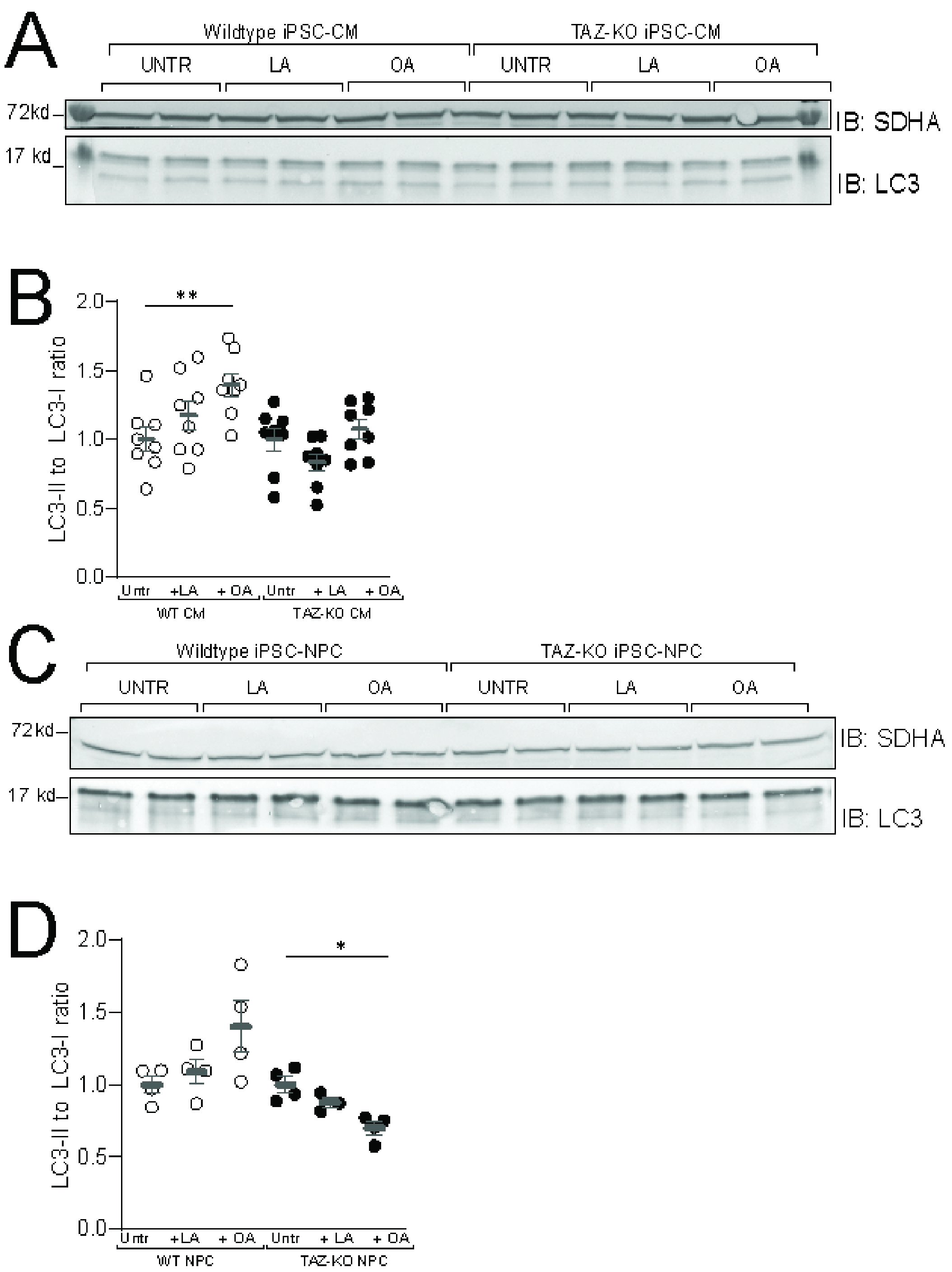


**Fig. S9.** ***Lipid treatment alters relative levels of autophagosome-associated LC3 in WT but not TAZ-KO CMs.*** (A and C) Whole cell lysate (60 µg) of indicated cell lines and treatments were immunoblotted for LC3-I (cytosolic) and LC3-II (lipidated, autophagosomal mebrane-associated). SDHA was used as a loading control. In (B) CMs and (D) NPCs, the ratio of LC3-II (16 kDa) to LC3-I (18 kDa) was calculated using ImageJ quantifiation of band intensities, normalized to loading control. LC3-II/-I ratio was normalized to the untreated control of each genotype. Significant differences are indicated; *≤0.05, **≤0.005, ***≤0.0005, and ****≤0.00005.

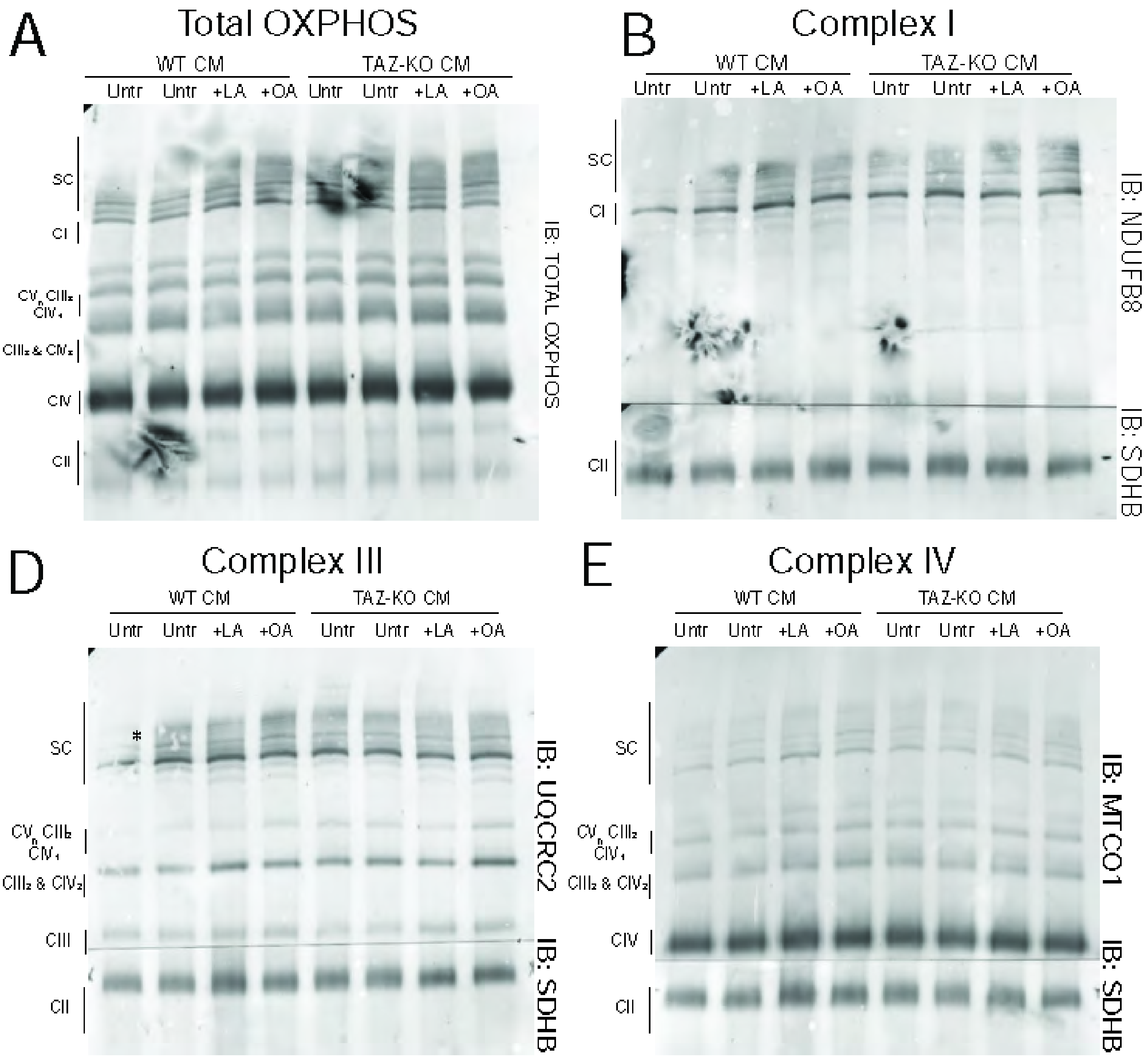


**Fig. S10.** ***Lipid treatment does not alter complex assembly in WT or TAZ-KO CMs.*** (A) Cell pellets of 1x10$^{6}$ cells of indicated cell type, genotype, and treatment, were resolved using blue native poly acrylamide gel electrophoresis (Blue Native- PAGE) and immunoblotting was performed to assess (A) total OXPHOS assembly (B) Complex I and II, (D) Complex III and II, and (E) Complex IV and II.

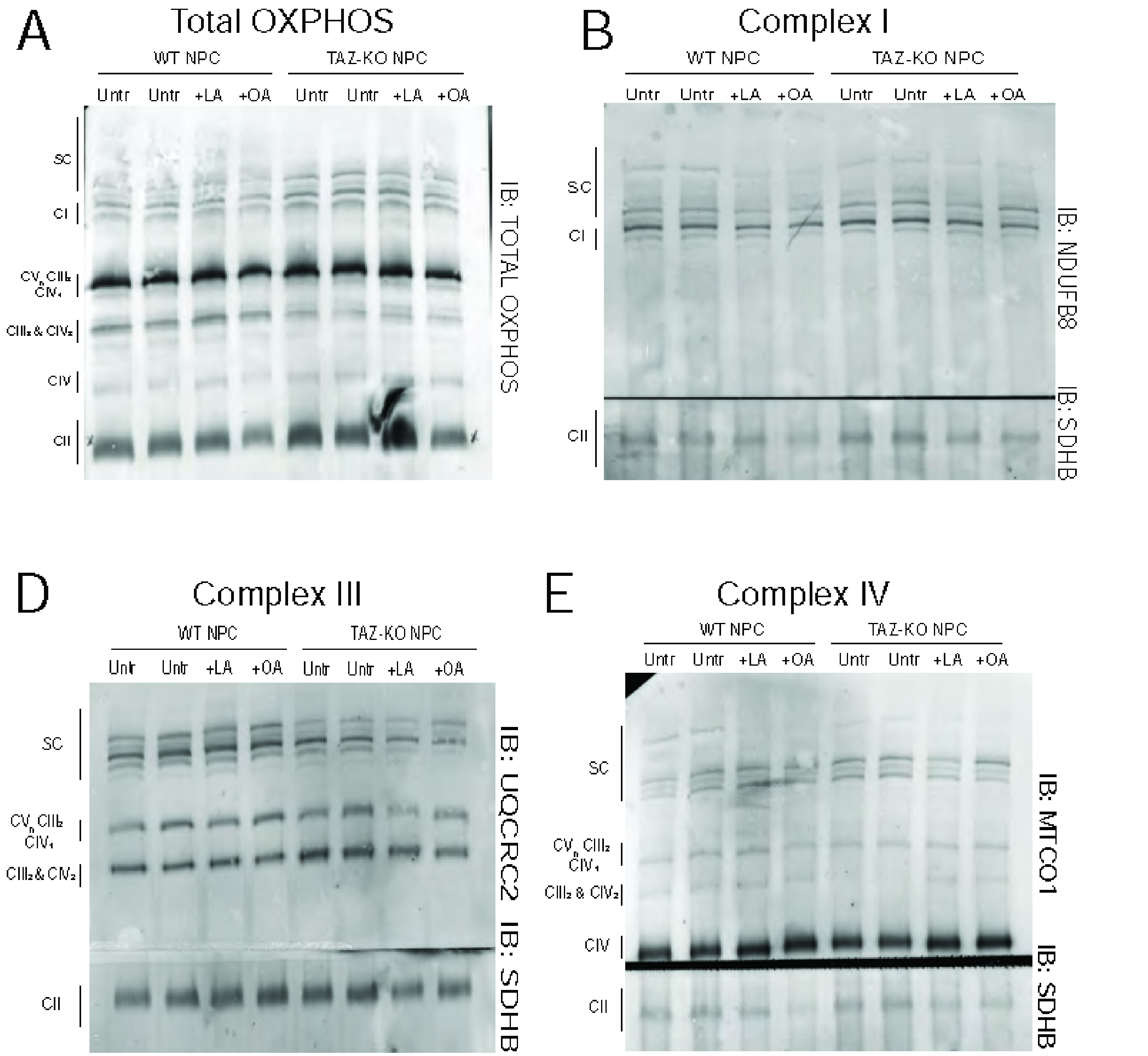


**Fig. S11.** ***Lipid treatment may rescue CIII-CIV defects in TAZ-deficient NPCs*** (A) Cell pellets of $1x10^6$ cells of indicated cell type, genotype, and treatment, were resolved using blue native poly acrylamide gel electrophoresis (Blue Native- PAGE) and immunoblotting was performed to assess (A) total OXPHOS assembly (B) Complex I and II, (D) Complex III and II, and (E) Complex IV and II. CIII-CIV intermediate complex reduction in TAZ-KO NPCs is partially remediated after 100µM LA or OA lipid supplementation.
